# Methylation of KRAS at lysine 182 and 184 by SETD7 promotes KRAS degradation

**DOI:** 10.1101/2021.02.10.429309

**Authors:** Chengyao Chiang, Tian Xiao, Songqing Fan, Hongmei Zheng, Shuaihu Li, Wenjun Guo, Min Zhang, Chuanqi Zhong, Juan Zeng, Duo Zheng

**Affiliations:** Guangdong Provincial Key Laboratory of Regional Immunity and Diseases, Shenzhen University International Cancer Center, Department of Cell Biology and Genetics, School of Medicine, Shenzhen University, Shenzhen, Guangdong, China; Department of Pathology, The Second Xiangya Hospital, Central South University, Changsha, Hunan, China; State Key Laboratory of Cellular Stress Biology, Innovation Center for Cell Signaling Network, School of Life Sciences, Xiamen University, Xiamen, Fujian, China; School of Biomedical Engineering, Guangdong Medical University, Dongguan, Guangdong, China

## Abstract

Oncogenic KRAS mutations are considered to be a key driver for initiation and progression in non-small-cell lung cancer (NSCLC). However, how post-translational modifications (PTMs) of KRAS, especially methylation, modify KRAS activity and downstream signals remain largely unclear. Here, we showed that SET domain containing histone lysine methyltransferase 7 (SETD7) interacts with KRAS and methylates KRAS at lysine 182 and 184. SETD7-mediated methylation of KRAS led to degradation of KRAS and attenuation of the RAS/MEK/ERK signaling cascade, endowing SETD7 with a potent tumor-suppressive role in NSCLC, both *in vitro* and *in vivo*. Mechanistically, RABGEF1, a ubiquitin E3 ligase of KRAS, was recruited and promoted KRAS degradation in a K182/K184 methylation-dependent manner. Notably, low SETD7 or RABGEF1 expression was associated with poor prognosis in lung adenocarcinoma patients. Taken together, our results establish a novel connection between lysine methylation and KRAS protein stability, in addition to elucidating a tumor-suppressive function of SETD7 that operates via modulation of oncogenic RAS signaling.

## Introduction

More than 30% of all human cancers are driven by mutations in RAS proto-oncogenes (Prior et al., 2012), which comprise four distinct isoforms: HRAS, NRAS, and two splice variants of the KRAS gene, KRAS-4A and KRAS-4B (Hobbs et al., 2016). The mutation frequencies of RAS isoforms differ in distinct cancer types. In pancreatic ductal adenocarcinoma (PDAC) and lung adenocarcinoma (LUAD), KRAS accounts for almost 100% of all RAS mutations (Cox et al., 2014). Oncogenic RAS mutations usually occur at the conserved residues G12, G13, or Q61 (Brose et al., 2002), which favor an active GTP-bound state and produce sustained activation of downstream signaling (Scheffzek et al., 1997; Trahey and McCormick, 1987). For their biological activity, RAS proteins must be anchored to the plasma membrane (PM) (Zhou et al., 2017), where they recruit a variety of effectors to transduce signals from tyrosine kinase receptors and stimulate signaling cascades (Simanshu et al., 2017).

Protein post-translational modifications (PTMs) are critical for translocation of RAS to the PM (Ahearn et al., 2018). Primary translation products of RAS undergo CAAX processing following a second signal; this signal comprises mono-palmitoylation on Cys residues located upstream of the CAAX motif for NRAS and KRAS-4A and duo-palmitoylation for HRAS (Hancock et al., 1989). KRAS-4B is not modified by palmitic acid (Hancock et al., 1989). Instead, it has a polybasic domain with eight lysines that are strongly positively charged and allow for electrostatic interactions with head groups of PM lipids (Hancock et al., 1990). Although KRAS-4B requires no second signal beyond CAAX processing to associate with membranes, the affinity of the interaction is still regulated by PTM. Phosphorylation of serine 181 (S181) by protein kinase C (PKC) within the polybasic region can partially neutralize the positive charge and mediate translocation of KRAS-4B from the PM to the endomembrane system (Bivona et al., 2006). S181 phosphorylation is essential for oncogenic KRAS function in activation of p-AKT and p-ERK1/2, as well as promoting cell proliferation, mobility, and tumor growth (Alvarez-Moya et al., 2010; Barcelo et al., 2014b). However, other types of conditional PTMs, especially methylation, and their role in modifying KRAS-4B localization and activity remain largely unknown.

Protein lysine methyltransferases (PKMTs) have been shown to methylate histone and non-histone proteins and participate in the regulation of several biological processes in both healthy human physiology and human diseases (Copeland et al., 2009). SET domain containing histone lysine methyltransferase 7 (SETD7), also known as SET7/9, KIAA1717, or KMT7, was first identified as a mono-methyltransferase of lysine 4 on histone H3 (H3K4) (Wang et al., 2001). The lysine residue methylated by SETD7 frequently followed the consensus amino acid motif [K/R]-[S/T/A](Del Rizzo and Trievel, 2011). Recently, several fundamental proteins involved in tumor progression have been identified as non-histone substrates of SETD7, including the tumor suppressor p53 (Campaner et al., 2011; Chuikov et al., 2004; Kurash et al., 2008), NF-κB p65 (Yang et al., 2009), hypoxia-induced factor-1α (HIF-1α) (Kim et al., 2016), β-catenin (Shen et al., 2015), and nuclear hormone estrogen receptor alpha (ERα) (Subramanian et al., 2008). The fates of proteins modified by SETD7-mediated lysine methylation are quite diverse. While methylation of p53 or ERα stabilized these proteins and enhanced transcriptional activity, methylation of β-catenin at K180 by SETD7 promoted ubiquitination and proteasomal degradation (Shen et al., 2015). Methylation at different lysines within a protein can even lead to divergent outcomes. For instance, SETD7 methylates the nuclear NF-κB p65 subunit at K37, which restricts p65 to the nucleus and facilitates its binding to promoters of inflammatory genes (Ea and Baltimore, 2009), whereas methylation at K314 and K315 destabilizes p65 by promoting ubiquitination (Yang et al., 2009). SETD7 can positively or negatively regulate multiple proteins; therefore, the function of SETD7 in tumor progression may be dependent on the cellular context and the substrate (Batista and Helguero, 2018). SETD7 contains three membrane occupation and recognition nexus (MORN) motifs in the N-terminal region, which is believed to mediate SETD7’s interaction with and anchorage to the PM (Bivona et al., 2006). However, to date, no proteins located in the PM have been identified as substrates of SETD7.

Here, by quantitative proteomics based on spectra count method, we determined that SETD7 interacts with KRAS on the PM and methylates KRAS at lysines 182 and 184. These PTMs promoted KRAS protein degradation by recruiting the E3 ligase, RabGEF1. Thus, SETD7 plays a potent tumor-suppressive role in KRAS-mutated NSCLC via the downregulation of RAS signaling. SETD7 was also downregulated in clinical lung cancer specimens and low SETD7 expression appeared to be associated with poor prognosis. Of note, SETD7 expression was inversely correlated with KRAS protein levels but not mRNA levels, further supporting the role played by SETD7 in PTMs of KRAS.

## Results

### SETD7 interacts with KRAS but not HRAS or NRAS

To identify the potential post-translational modifications of KRAS-4B (hereafter referred to as KRAS), we first explored KRAS interactome in the human embryonic kidney 293T cells *via* the ectopic expression of 3×HA-tagged KRAS (HA-KRAS). We prepared cellular extracts and incubated them with anti-HA magnetic beads. After extensive washing, bound proteins were eluted with a basic buffer and resolved by SDS-PAGE (**Figure 1A**), and then followed by liquid chromatography tandem mass spectrometry (LC-MS/MS) analysis (**Figure 1B**). As expected, Ras GTPase-activating protein-binding protein 1 and 2 (G3BP1/G3BP2), as well as Ras-related proteins RAB11A/RAB7A/RAB6A were detected as candidate KRAS-interacting proteins (**Supplementary Table 1**). Consistent with previous report, ribonucleoprotein HNRNPA2B1 which interacted with and regulated oncogenic KRAS in PDAC cells(Barcelo et al., 2014a) was also found in the KRAS interactome. Of note, by quantitative proteomics based on spectra count method, SETD7 was identified as the most significantly differential protein enriched in the immunoprecipitation (IP) sample of HA-KRAS-expression cells compared to that of vector control (**Figure 1C, Supplementary Figure S1A**). Intriguedly, SETD7 only interacted with KRAS and not the other two RAS family members, HRAS and NRAS (**Figure 1D**). Confocal microscopy analysis showed that SETD7 was co-localized with KRAS, predominantly on the plasma membrane, where KRAS recruits regulators and effectors to exert its function (**Figure 1E**). The interaction between KRAS and SETD7 was further confirmed by co-immunoprecipitation (co-IP) in both directions in the NSCLC cell line NCI-H358 (H358) with stable SETD7 overexpression (**Figure 1F and G**). Endogenous KRAS protein was also immunoprecipitated from A549 cell extracts using anti-SETD7 antibody, indicating an interaction between KRAS and SETD7 at the physiological level (**Figure 1H**).

**Figure 1.**
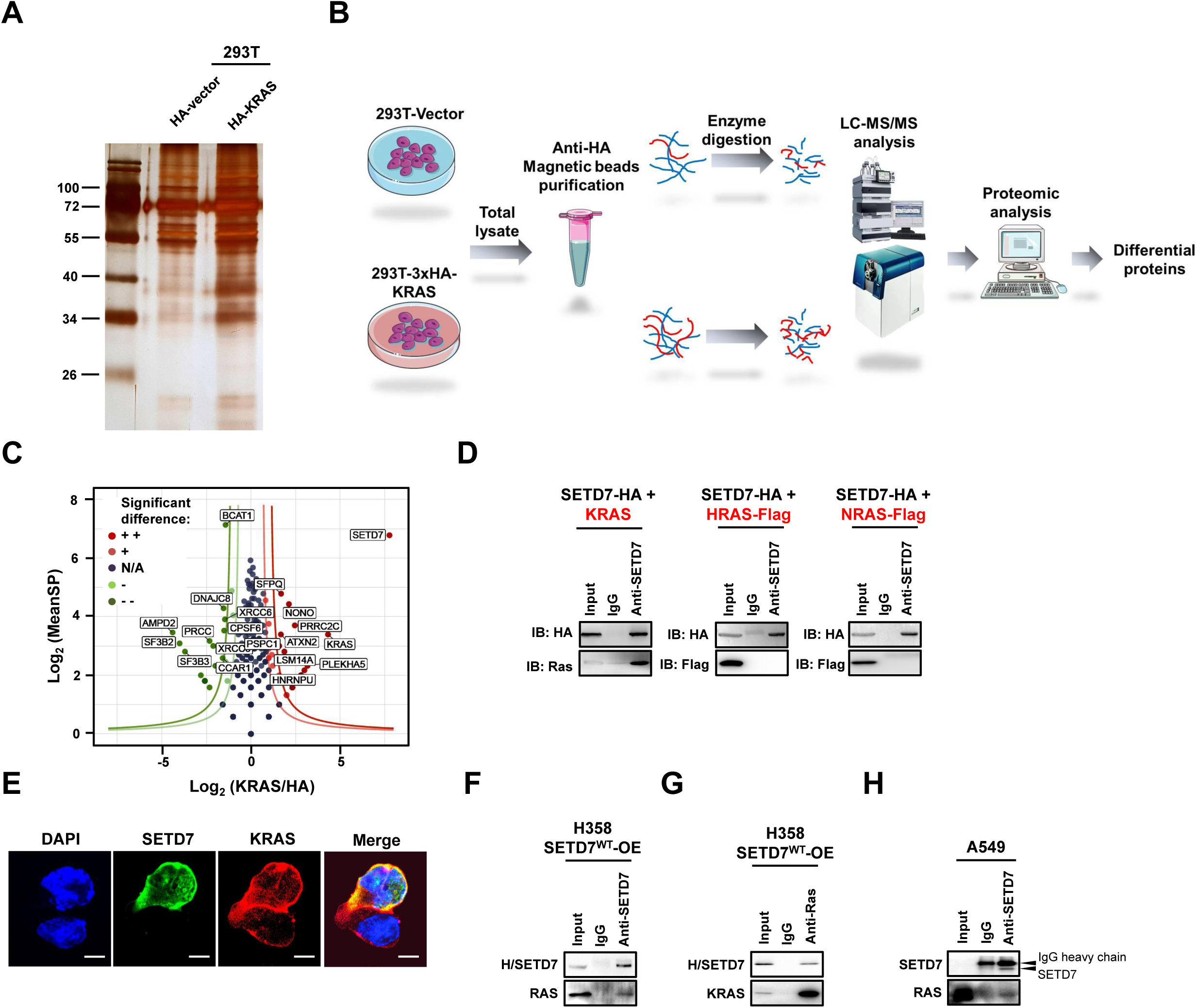
SETD7 interacts with KRAS. (A) Proteins from the co-immunoprecipitation (co-IP) assay using anti-HA magnetic beads in 293T with ectopic expression of 3×HA-tagged KRAS were separated by polyacrylamide gel electrophoresis and visualized using silver staining. **(B)** The workflow for identifying KRAS-interacting proteins was shown, including co-IP assay, sample preparation, LC-MS/MS analysis, protein quantification, and further computational analyses for differential proteins. **(C)** Volcano plot of proteins identified by LC-MS/MS analysis in HA-vector and 3×HA-KRAS groups. The ratio (KRAS/HA) and the mean of protein spectra counts (MeanSP) in two groups were calculated. The x axis shows the log2 ratio (KRAS/HA) and the y axis shows the log2 MeanSP. The proteins distributed outside the dark-red boundary line (y = log_2_2.5 / (x-1)) were labeled as significant differential proteins in KRAS group. TOP 10 differential proteins from each group were labeled. **(D)** Western blotting (WB) analysis of whole-cell lysates and immunoprecipitation (IP) of SETD7 in 293T cells transfected with the indicated plasmids. **(E)** Immunofluorescence staining of SETD7 (green) and KRAS (red) in 293T cells co-transfected with SETD7 and KRAS. DAPI was used to label the nuclei. Scale bar, 10 µm. **(F and G)** WB analysis of the co-IP assay using anti-SETD7 (F) or anti-RAS (G) antibodies in H358-SETD7-O/E stable cells. (**H**) WB analysis of the co-IP assay with anti-SETD7 antibody in non-small-cell lung cancer (NSCLC) cell line A549.

### SETD7 is a potential tumor suppressor in lung cancer cells with KRAS mutations

To understand the role of SETD7 in NSCLC progression, we first checked the expression of SETD7 in NSCLC cell lines and clinical specimens. According to staging information relating to NSCLC cell lines, obtained from the American Type Culture Collection (ATCC), we found that SETD7 expression was much lower in cell lines derived from late-stage tumors (**Supplementary Figure S1B**).

Since SETD7 exclusively interacts with KRAS, and not HRAS and NRAS, we next examined the potential function of SETD7 in NSCLC cell lines harboring KRAS mutations. In H358 (KRAS G12C) and A549 (KRAS G12S) cells, overexpression (OE) of SETD7 impaired both cell proliferation and two-dimensional (2-D) colony formation (**Figure 2A and B, Supplementary Figure S2A and B**). The spheroid growth of H358 cells in soft-agar culture was also attenuated with ectopic SETD7 OE (**Figure 2C**), implying a tumor-suppressive effect of SETD7 in NSCLC. To test whether the anti-tumor effect of SETD7 was associated with its interaction with KRAS, we examined the impact of SETD7 on KRAS and downstream signaling cascades. Interestingly, along with downregulation of KRAS at the protein level, two critical downstream effectors, phospho-ERK1/2 (p-ERK1/2) and phospho-AKT (p-AKT), also apparently decreased when SETD7 was overexpressed in H358 (**Figure 2D**) and A549 cells (**Supplementary Figure S2C**). On the other hand, efficient silencing of SETD7 by short hairpin RNAs (shRNAs) strongly promoted cell growth and colonization (**Figure 2E and F, Supplementary Figure S2D and E**), in addition to enhancing anchorage-independent cell growth in NCI-H1437 (H1437) (**Figure 2G, Supplementary Figure S2F)** and H358 cells (**Supplementary Figure S2G**). As expected, knockdown (KD) of SETD7 abrogated its inhibition of KRAS expression, leading to elevated levels of KRAS protein and the reactivation of p-ERK1/2 and p-AKT (**Figure 2H and Supplementary Figure S2H**).

**Figure 2.**
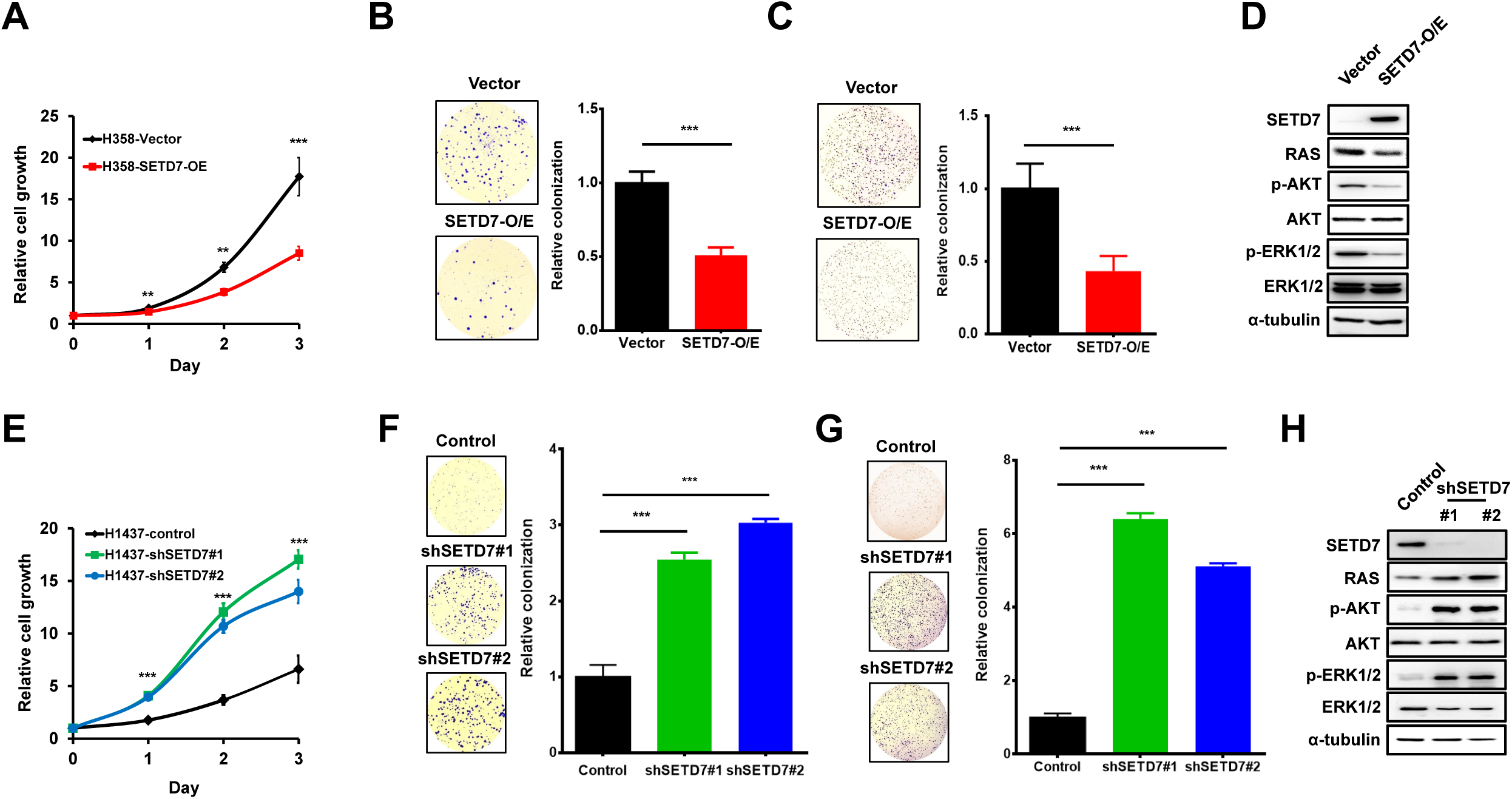
SETD7 plays a tumor-suppressive role in non-small-cell lung cancer (NSCLC) cell lines by downregulating RAS-associated signaling. **(A-C)** Cell proliferation (A), two-dimensional (2-D) colony formation (B), and anchorage-independent cell growth (C) of H358-vector and H358-SETD7-O/E stable cells were measured separately. **(D)** Western blotting (WB) analysis of SETD7 and RAS-related signaling pathways in H358 stable cells. **(E-G)** Cell proliferation (E), 2-D colony formation (F), and anchorage-independent cell growth (G) of H1437-control and two SETD7-KD stable cells were measured separately. The bar graphs represent the quantification of numbers of colonies on plates or spheres in soft agar from three independent experiments. **(H)** WB analysis of SETD7 and RAS-related signaling pathways in H1437 stable cells. \*\**P*<0.01; \*\*\**P*<0.001.

We further investigated the anti-tumor effect of SETD7 *in vivo*. A549 cells with stable expression of SETD7 or a vector control were subcutaneously injected into the opposite flanks of nude mice. The growth rate of xenograft tumors with SETD7 OE was significantly attenuated (**Figure 3A**). Compared with the results of paired control tumors, both the volume and weight of tumors with SETD7 OE were lower at 7 weeks post-injection (**Figure 3B, Supplementary Figure S3A**). Immunohistochemical staining of tumor tissues revealed that SETD7 downregulated endogenous KRAS expression and inhibited tumor cell proliferation, which was characterized by a decreased percentage of Ki67-positive cells (**Figure 3C**). For H1437 cells with relatively high SETD7 expression (**Supplementary Figure S1B**), their tumorigenic ability in nude mice seemed poor, as growth of the subcutaneous xenograft was very slow and even ceased one week post-injection (**Figure 3D**). Notably, SETD7 knockdown in H1437 cells greatly accelerated xenograft tumor growth (**Figure 3D**) and consequently boosted both the volume and weight of tumors with SETD7 KD at 4 weeks post-injection **(Figure 3E, Supplementary Figure S3B)**. As a consequence of SETD7 knockdown, both the KRAS level and percentage of Ki67-positive cells were simultaneously elevated in xenograft tumor tissues (**Figure 3F**). Taken together, these *in vitro* and *in vivo* findings indicate that SETD7 suppressed NSCLC progression via the negative regulation of KRAS expression and downstream MEK/ERK and PI3K/AKT signaling pathways.

**Figure 3.**
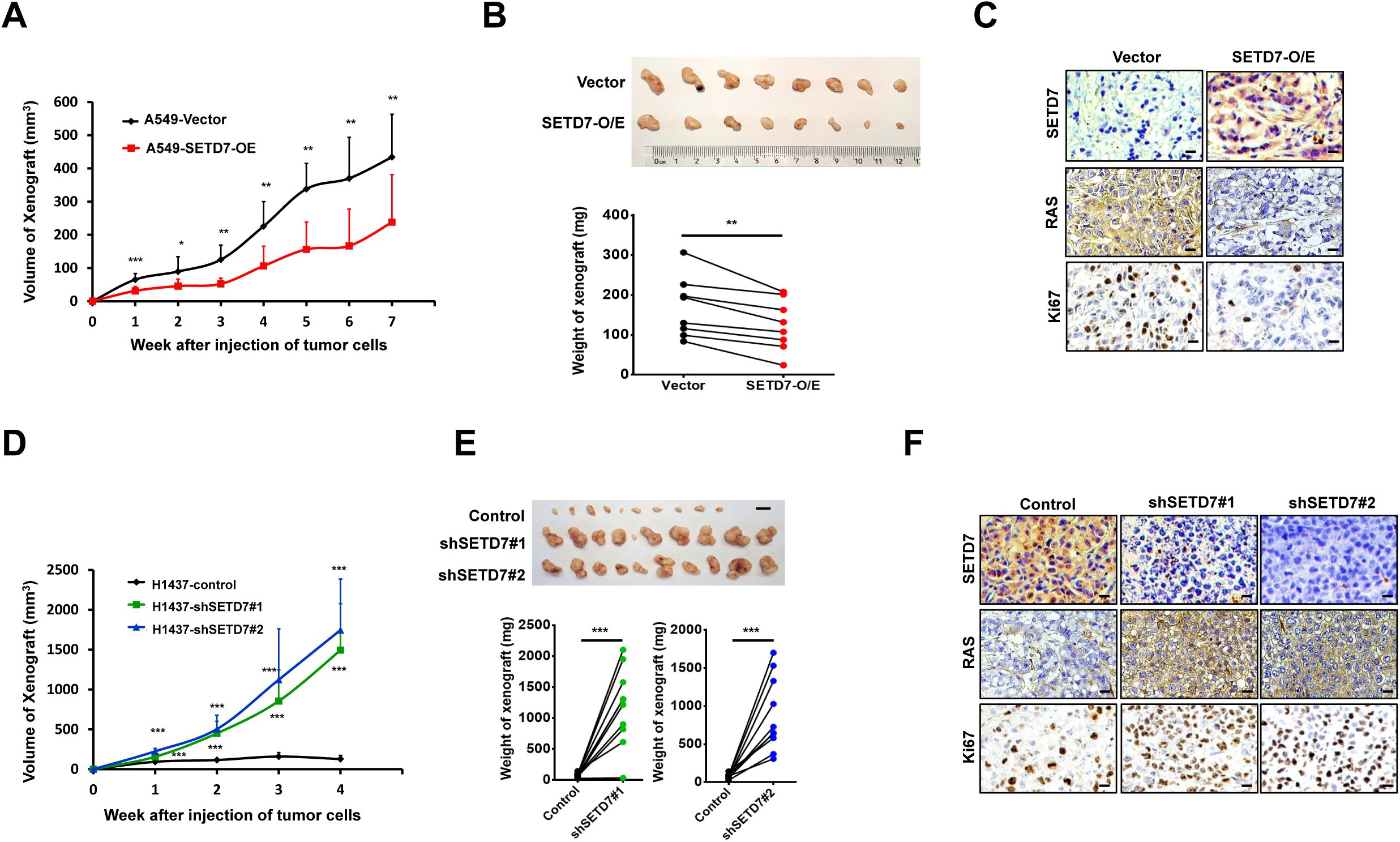
The anti-cancer effect of SETD7 *in vivo*. **(A)** Measurement of volumes of xenograft tumors derived from A549-vector or A549-SETD7-O/E stable cells (n = 8). **(B)** A photograph of the xenograft tumors and the tumor weights, 7 weeks after cell injection, are shown (upper and lower panels, respectively). **(C)** Representative images of immunohistochemical (IHC) staining of SETD7, RAS, and Ki67 in xenograft tumor tissues. Scale bar, 100 μm. **(D)** Measurement of volumes of xenograft tumors derived from H1437-control and two SETD7-KD stable cells (n = 10). **(E)** A photograph of the xenograft tumors and the tumor weights, 4 weeks after cell injection, are shown (upper and lower panels, respectively; scale bar, 10 mm). **(F)** Representative images of IHC staining of SETD7, RAS, and Ki67 in xenograft tumor tissues of each group. Scale bar, 100 μm. \**P*<0.05; \*\**P*<0.01; \*\*\**P*<0.001.

### SETD7 promotes KRAS ubiquitination and degradation

To elucidate the molecular mechanism underlying SETD7-mediated KRAS downregulation, we first sought to determine whether SETD7 impinges on KRAS transcription. Quantitative PCR results showed that neither SETD7 OE in A549 and H358 cells (**Supplementary Figure S4A and B**) nor knockdown in H1437 cells changed KRAS mRNA expression (**Supplementary Figure S4C**). Thus, we speculated that the interaction of SETD7 with KRAS may influence KRAS protein stability. To test this hypothesis, the protein levels of KRAS were determined at different time points following treatment with cycloheximide (CHX), an inhibitor of protein synthesis in eukaryotes. We found that co-expression of SETD7 and KRAS in 293T cells accelerated the degradation of exogenous KRAS proteins (**Figure 4A**). Additionally, SETD7 OE obviously stimulated the turnover rate of endogenous KRAS in H358 and A549 cells (**Figure 4B and Supplementary Figure S4D**), whereas knockdown of SETD7 prolonged the half-life of KRAS proteins (**Figure 4C**).

**Figure 4.**
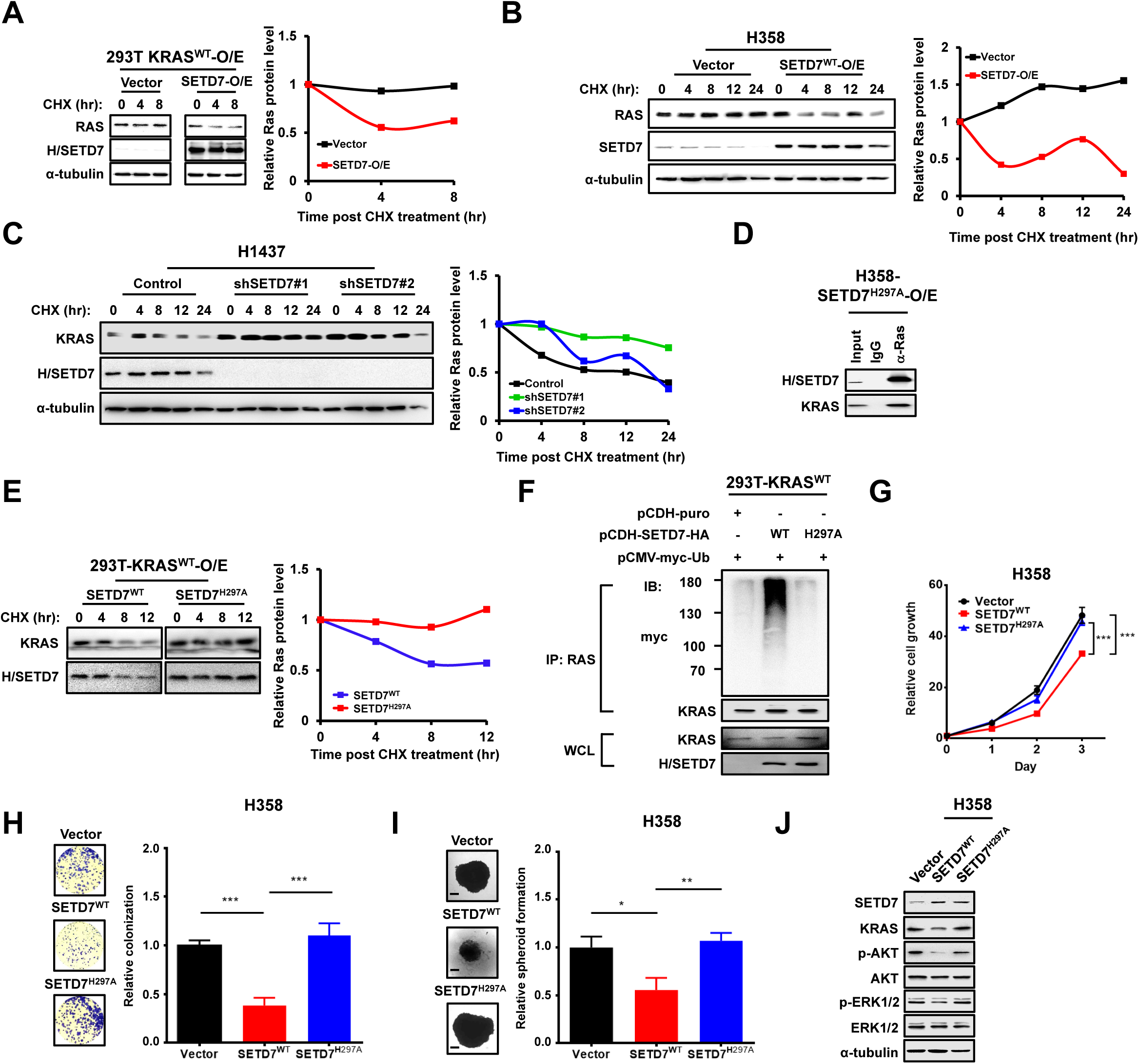
SETD7 promotes KRAS degradation and ubiquitination in a catalytic-dependent manner. **(A)** Western blotting (WB) analysis of 293T cells transfected with KRAS and SETD7 or KRAS alone with cycloheximide (CHX) (100 μg/mL) treatment, as indicated. **(B)** WB analysis of H358-vector and H358-SETD7-O/E stable cells with CHX (100 μg/mL) treatment, as indicated. **(C)** WB analysis of H1437-control and two SETD7-KD stable cells with CHX (100 μg/mL) treatment, as indicated. The graphs in A-C represent the quantification of RAS protein levels relative to the α-tubulin level. **(D)** WB analysis of the co-immunoprecipitation (co-IP) assay using anti-RAS antibody in H358 cells stably expressing the SETD7 H297A mutant. **(E)** WB analysis of 293T cells co-transfected with KRAS and wild-type SETD7 or SETD7 H297A mutant with CHX (100 μg/mL) treatment, as indicated. The graph represents the quantification of RAS protein levels relative to the α-tubulin level. **(F)** 293T cells transfected with the indicated combinations of KRAS, HA-SETD7 (WT or H297A), and myc-Ubi were subjected to a ubiquitination assay. **(G-I)** Cell proliferation (G), 2-D colony formation (H) and spheroid growth (I) of H358-vector, H358-SETD7^WT^, and H358-SETD7^H297A^ stable cells were measured separately. \**P*<0.05; \*\**P*<0.01; \*\*\**P*<0.001. **(J)** WB analysis of SETD7 and RAS-related signaling pathways in H358 stable cells, as indicated.

To determine whether SETD7 methyltransferase activity is indispensable for KRAS degradation and ubiquitination, we constructed a catalytically dead SETD7^H297A^ mutant(Nishioka et al., 2002). As expected, OE of wild-type (WT) SETD7 markedly enhanced degradation (**Figure 4E**) and ubiquitination of KRAS (**Figure 4F**). However, the H297A mutant of SETD7, which lacks lysine methyltransferase activity(Nishioka et al., 2002), could interact with KRAS (**Figure 4D**) but had lost the ability to promote KRAS degradation (**Figure 4E**) and ubiquitination (**Figure 4F**), suggesting this catalytic activity is critical for SETD7-mediated KRAS ubiquitination and degradation. The suppression of cell growth and 2-D colony formation by WT SETD7 in H358 and A549 cells was reversed by the H297A mutant (**Figure 4G and H, Supplementary Figure S4E and F**). SETD7^WT^-inhibited growth of three-dimensional (3-D) spheroids in H358 and A549 cells was also restored by the H297A mutant (**Figure 4I and Supplementary Figure S4G**). Accordingly, SETD7^WT^ repression of the KRAS-related pathway was rescued by SETD7^H297A^ (**Figure 4J and Supplementary Figure S4H**). These results imply that SETD7 reduces KRAS protein stability through ubiquitination, in a catalytic-dependent manner.

### SETD7 methylates KRAS at K182 and K184 and facilitates E3 ligase RabGEF1 recruitment

We determined that SETD7-mediated KRAS ubiquitination and degradation were dependent on its methyltransferase activity, so we further investigated whether KRAS is a direct substrate of SETD7. Among known SETD7 substrates, the methylated lysine site usually follows the conserved amino acid motif [K/R]-[S/T/A](Del Rizzo and Trievel, 2011). By analyzing the amino acid sequence of KRAS-4B, two lysine residues close to the C-terminal, lysine 182 (K182) and lysine 184 (K184), were identified as potential target methylation sites for SETD7 **(Figure 5A)**. However, the consensus motif was not found in KRAS-4A, NRAS, or HRAS (data not shown). Using a computer simulation, the complex between SETD7 and KRAS could only be stabilized in trajectory from structure (str.) 4 of the simulated styles (**Supplementary Figure S5A and B**). Based on the str. 4 model, K182 and K184, together with T183, formed an interactional network with SETD7 and helped KRAS bind to SETD7 (**Figure 5B**). As predicted, KRAS methylation was enhanced by co-expression of WT SETD7 but not the catalytically dead H297A mutant (**Figure 5C**). Mutation of K182 or K184 to methionine obviously reduced the level of KRAS methylation compared with that of the WT group (**Figure 5D**). Of note, the interaction between SETD7 and KRAS was also attenuated following K182 and K184 mutation (**Figure 5D**). To further confirm SETD7 methylates KRAS at K182 and K184, we performed immunoprecipitation with anti-HA magnetic beads in 293T cells with co-expression of SETD7^WT^ (or SETD7^H297A^) and 3×HA-tagged KRAS, and identified the lysines methylation on KRAS using LC-MS/MS analysis. K182 methylation of KRAS was detected in SETD7^WT^ group but not in SETD7^H297A^ group, supporting our novel finding that KRAS is a direct substrate of SETD7 (**Figure 5E and Supplementary Figure S5C**).

**Figure 5.**
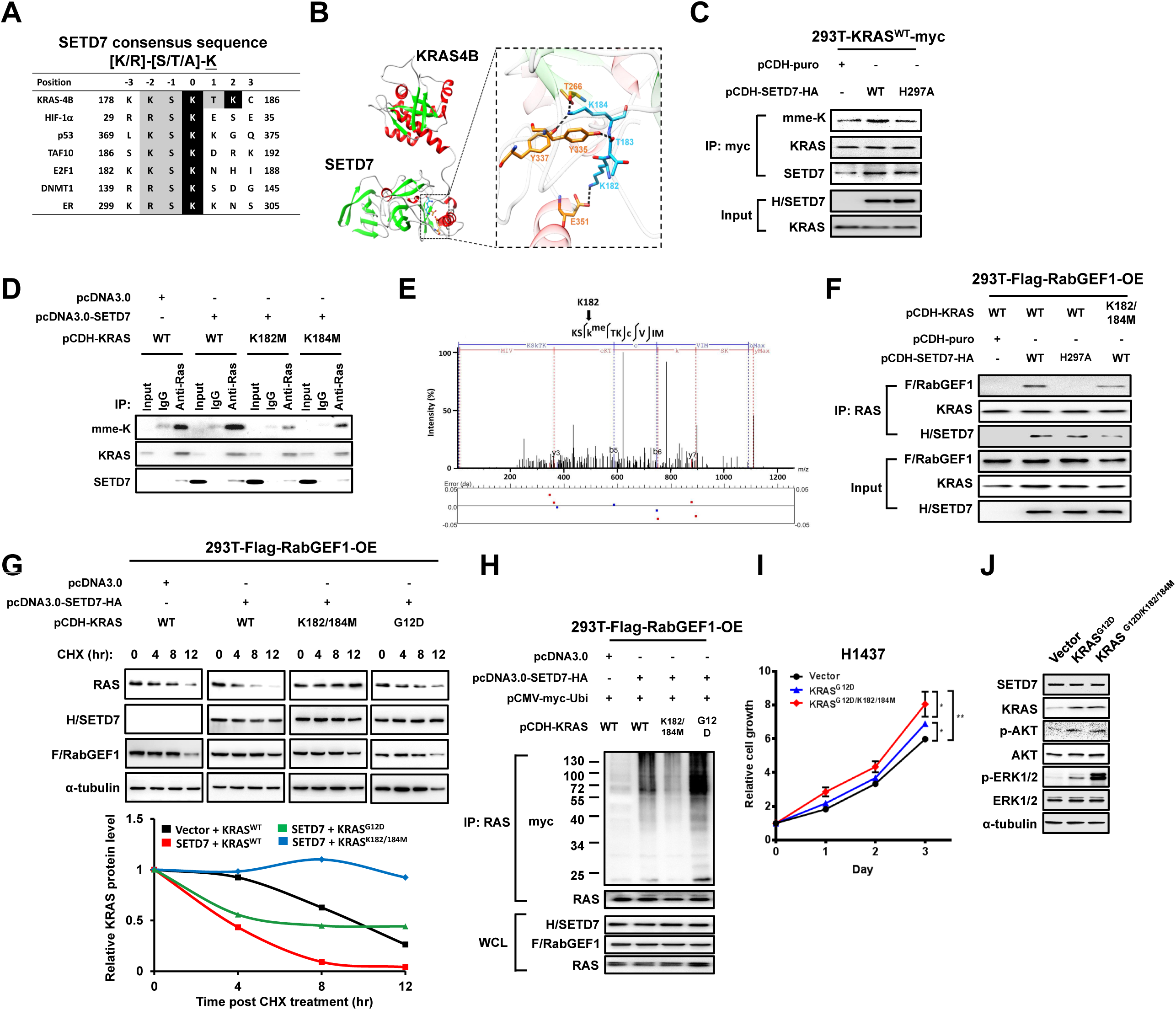
SETD7 methylates KRAS at K182 and K184, leading to KRAS ubiquitination and degradation via recruitment of RabGEF1. (**A**) Alignment of amino acid sequences from proteins reported to be SETD7 substrates and amino acids 178-186 in the C-terminal of KRAS. Lysines 182 and 184 of KRAS were predicted to be potential residues for SETD7-mediated methylation. (**B**) A representative structure of the interactional network between SETD7 and KRAS, based on a cluster analysis. K182, T183, and K184 of KRAS and some important residues of SETD7 are represented by a stick model. Residues from SETD7 and KRAS are colored orange and cyan, respectively. (**C**) 293T cells were co-transfected with KRAS-myc and wild-type SETD7 or SETD7 H297A mutant. KRAS was immunoprecipitated (IP) using anti-myc antibody and the methylation level of KRAS was detected using mono-methyl lysine (mme-K) antibody. (**D**) KRAS methylation was detected in 293T cells transfected with the indicated plasmids and IP using anti-RAS antibody. (**E**) KRAS peptide modified with methylation on lysine 182 residue was identified by LC-MS/MS. (**F**) Western blotting (WB) analysis of whole-cell lysates (WCLs) and IP of RAS in 293T cells transfected with the indicated plasmids. (**G**) WB analysis of 293T cells transfected with combinations of RabGEF1, HA-SETD7, and KRAS (WT, K182/184M, or G12D) with CHX (100 μg/mL) treatment, as indicated. The graph represents the quantification of RAS protein levels relative to the α-tubulin level. (**H**) 293T cells transfected with the indicated combinations of RabGEF1, HA-SETD7, KRAS (WT, K182/184M, or G12D), and myc-Ubi were subjected to a ubiquitination assay. **(I)** Cell proliferation of H1437-vector, H1437-KRAS^G12D^, and H1437-KRAS^G12D/K182/184M^ stable cells was measured using a CCK-8 assay. \**P*<0.05; \*\**P*<0.01. **(J)** WB analysis of SETD7 and RAS-related signaling pathways in H1437 stable cells, as indicated.

We next sought to investigate the underlying mechanism via which methylation at K182 and K184 regulates KRAS protein stability. It has been reported that RabGEF1 is an E3 ligase of the RAS family members HRAS and NRAS(Xu et al., 2010). Thus, we asked whether RabGEF1 could act as an E3 ligase of KRAS. A co-IP assay showed an interaction between KRAS and RabGEF1 only in the presence of SETD7^WT^ but not in SETD7^H297A^ group (**Figure 5F**). In addition, dual mutations of KRAS at K182 and K184 (KRAS-DM) restricted its interaction with RabGEF1, suggesting that methylation of KRAS by SETD7 facilitates the recruitment of RabGEF1 (**Figure 5F**). While RabGEF1 alone could not promote KRAS degradation (**Supplementary Figure S5D**), the co-expression of SETD7 and RabGEF1 worked synergistically to reduce the half-lives of KRAS proteins (**Supplementary Figure S5E**). In contrast, KRAS-DM exhibited greater protein stability, which was impervious to SETD7/RabGEF1-mediated degradation (**Figure 5G**). Moreover, knockdown endogenous RabGEF1 increased stability and extended the half-lives of KRAS proteins (**Supplementary Figure S5F**). Consistent with previous study(Xu et al., 2010), RabGEF1 alone couldn’t induce KRAS ubiquitination (**Figure 5H**). However, in accordance with the reduced half-life of KRAS, SETD7 and RabGEF1 cooperated to increase the ubiquitination of WT KRAS, whereas KRAS-DM exhibited a ubiquitination-resistant phenotype (**Figure 5H**). These results suggest that RabGEF1-induced KRAS degradation and ubiquitination is dependent on SETD7-mediated K182/184 methylation of KRAS protein.

Interestingly, in the presence of both SETD7 and RabGEF1, the protein stability of the constitutively active KRAS G12D mutant also decreased (**Figure 5G**), accompanying with the increase of ubiquitination (**Figure 5H**), implying that SETD7/RabGEF1-mediated KRAS degradation may be widespread in mammalian cells, regardless of KRAS mutation status. Compared with KRAS G12D activity, KRAS G12D-DM showed a more obvious effect on promoting cell proliferation (**Figure 5I)** and activating downstream MEK/ERK and PI3K/AKT signaling in H1437 cells (**Figure 5J**), suggesting that the prolonged half-life and increased protein stability conferred a more potent function on KRAS. Taken together, these data indicate that methylation of KRAS by SETD7 at the K182 and K184 residues promotes ubiquitination and degradation through the recruitment of the E3 ligase RabGEF1.

### SETD7 is downregulated in clinical NSCLC samples and inversely correlated with KRAS at the protein level

Since SETD7 suppressed NSCLC progression by promoting KRAS methylation and degradation, we further sought to determine the clinical significance of SETD7 in NSCLC. Analysis of the Kaplan-Meier (KM) plotter database (Gyorffy et al., 2013) revealed that increased SETD7 expression was associated with better prognosis in patients with NSCLC or lung adenocarcinoma (LUAD) (**Figure 6A)**, but not in patients with lung squamous cell carcinoma (LUSC) **(Supplementary Figure S6A**). Our database analysis also showed that RabGEF1 acted as a tumor suppressor in lung cancer, especially LUAD (**Supplementary Figure S6B**). The combined SETD7 and RabGEF1 profiles showed that only patients with simultaneously increased expression of SETD7 and RabGEF1 exhibited longer survival times **(Supplementary Figure S6C)**. In the population with increased SETD7 expression, decreased RabGEF1 expression was clearly associated with poor survival in LUAD, and vice versa (**Supplementary Figure S6D and E)**. More importantly, in the decreased SETD7-expression (or RabGEF1) population, increased expression of RabGEF1 (or SETD7) did not confer any survival advantage, implying that SETD7 and RabGEF1 might cooperate to execute their tumor-suppressive roles in LUAD progression (**Supplementary Figure S6F and G**). In our own clinical cohort with NSCLC, the expression of SETD7 varied among patients (**Figure 6B)** and those who possessed high SETD7 expression also displayed better prognoses (**Figure 6C**).

**Figure 6.**
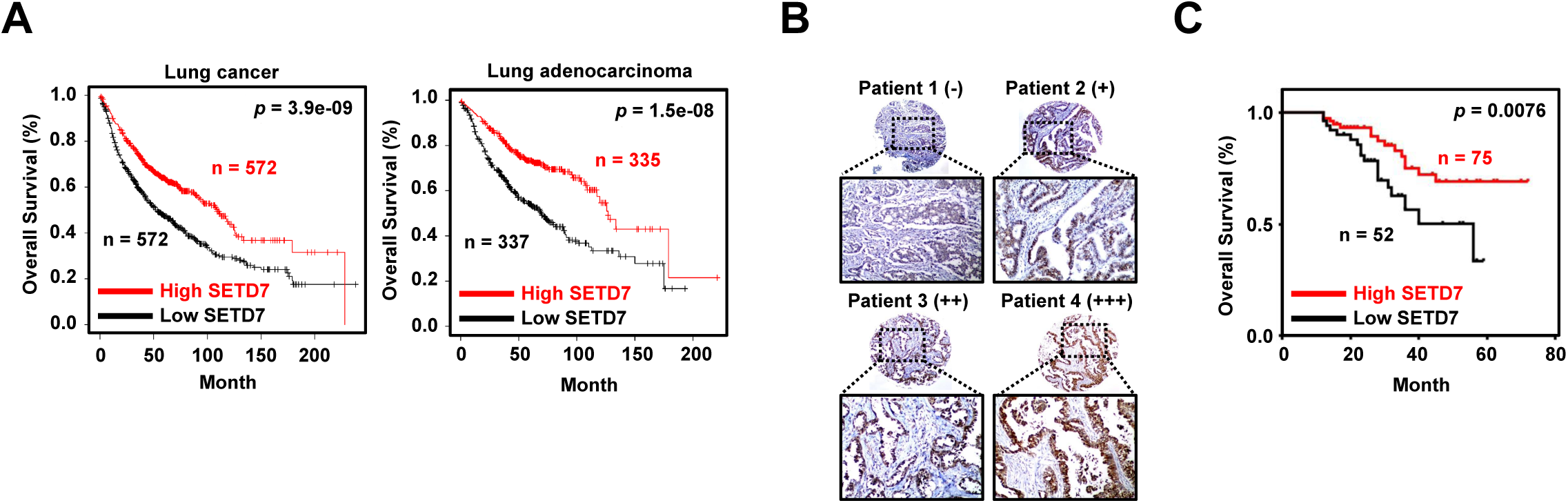
The clinical significance of SETD7 in lung adenocarcinoma (LUAD) (**A**) Kaplan-Meier plotter survival analysis for SETD7 mRNA levels (probe set, 224928_at) in lung cancer (n = 1144) and LUAD (n = 672) patient samples. Medium was used as a cutoff to divide the samples into low and high groups. **(B)** Representative images of immunohistochemical (IHC) staining of SETD7 in a cohort of 127 LUAD tissue samples. The staining intensity was classified as negative (-), weak (+), moderate (++), or strong (+++). **(C)** Survival analysis for IHC staining intensity of SETD7 in LUAD tissue samples. Negative and weak staining was indicated as SETD7-low (n = 52), while moderate and strong staining was indicated as SETD7-high (n = 75).

## Discussion

Here, we describe for the first time that lysine residues on KRAS protein are methylated by SETD7. SETD7 interacts with KRAS and methylates it at K182 and K184. This methylation destabilizes KRAS via the enhancement of ubiquitin-mediated protein degradation by the E3 ligase RabGEF1. Interestingly, the epigenetic codes written by SETD7 appear critical for the recruitment of RabGEF1 to KRAS. KM plotter analysis also showed SETD7 and RabGEF1 act synergistically to prolong overall survival in patients with LUAD. These findings imply that SETD7 may cooperate with RabGEF1 to exert an anti-cancer effect in NSCLC, and that the combination of SETD7 and RabGEF1 may serve as a useful index for prognostic prediction in KRAS-driven LUAD.

HRAS, NRAS, KRAS4A, and KRAS4B share more than 80% sequence identity in their N-terminals. The variation in the C-terminal among the four isoforms is critical for determining the subcellular localization of RAS (Simanshu et al., 2017). Although phosphorylation of serine 181 (S181) on the C-terminal tail of KRAS-4B by protein kinase C (PKC) has been well studied for its role in regulating KRAS translocation and signaling transduction, the outcome of KRAS phosphorylation at S181 remains controversial. Some studies have shown that S181 phosphorylation of KRAS, which is inhibited by calmodulin (CaM), is essential for oncogenic KRAS function in activation of p-AKT and p-ERK1/2, as well as promoting cell proliferation, mobility, and tumor growth (Alvarez-Moya et al., 2010; Barcelo et al., 2014b). However, others have reported that PKC-mediated S181 phosphorylation could either induce KRAS-4B translocation from the PM to mitochondria and trigger Bcl-XL-dependent apoptosis (Bivona et al., 2006), or impair the KRAS-CaM interaction and activate non-canonical Wnt signaling, which prevents tumorigenicity (Wang et al., 2015). Specifically, phosphorylation of GTP-bound KRAS-4B on S181 was shown to reduce nanocluster formation (Plowman et al., 2008), a spatial organization of RAS upon which signal transmission is dependent (Tian et al., 2007). A negatively charged phosphate group on S181 would repel cell membrane phospholipids, trigger the dissociation of KRAS from the cell membrane, and transport KRAS back to subcellular compartments (Feig, 2006); therefore, it is most likely that S181 phosphorylation negatively regulates KRAS function. Thus, PKC agonists or activators have been proposed for use as anti-cancer agents to inhibit oncogenic, KRAS-driven malignancies (Bivona et al., 2006; Wang et al., 2015). In our study, we found that SETD7 overexpression also induced translocation of KRAS from the PM to the cytosol, similar to the effect of S181 phosphorylation, whereas K182/K184 mutations could reverse this phenotype (unpublished data). Since methylation does not change the positive charge of lysine residues, we speculated that K182/K184 methylation might be a prerequisite for S181 phosphorylation of KRAS. We further noticed that K182/184M or S181A mutations of KRAS were resistant to ubiquitination-mediated degradation (Figure 6H and unpublished data), suggesting that PTM of residues located in polybasic domains links the subcellular localization of KRAS to its protein stability. Thus, further elucidation of the interplay between K182/K184 methylation and S181 phosphorylation will be important for understanding how different types of PTM cooperate to modulate KRAS protein stability.

SETD7 is an important modifier of several non-histone proteins and exerts either oncogenic or tumor-suppressive roles in different types of cancer, depending on the cellular context (Batista and Helguero, 2018). Previous studies have shown that SETD7 serves as a tumor suppressor in breast cancer (Montenegro et al., 2016), renal cellular carcinoma (Liu et al., 2015), colorectal cancer, and gastric cancers(Hong et al., 2018). Nevertheless, an oncogenic role for SETD7 has also been reported in breast cancer (Subramanian et al., 2008), prostate cancer (Wang et al., 2018), hepatocellular carcinoma (Chen et al., 2016), and intestinal tumorigenesis (Oudhoff et al., 2016), via various molecular mechanisms. The controversial role SETD7 plays in different types of cancer suggests that the function of SETD7 in cancer progression depends on the cellular or tissue context and specific interacting proteins. We found that SETD7 inhibited KRAS-driven LUAD both *in vitro* and *in vivo*. Surprisingly, in NSCLC cell lines with constitutively active EGFR mutations, SETD7 overexpression promoted cell growth (unpublished data). Since EGFR mutant cell lines were not sensitive to inhibition of MEK (Tricker et al., 2015), a direct downstream effector of RAS, SETD7 may interact with other, undefined substrates and play an oncogenic role in an EGFR-mutated genetic context. To date, KRAS-driven lung cancer remains an intractable disease. In view of the promising anti-cancer effect of SETD7 in NSCLC cell lines with KRAS addiction, approaches involving the upregulation of SETD7 expression or activation of SETD7 enzymatic activity might benefit lung cancer patients with KRAS mutations. Furthermore, since KRAS and EGFR mutations are usually mutually exclusive in LUAD (Ding et al., 2008), further exploration of the distinct roles of SEDT7 in KRAS- or EGFR-mutant contexts, which are the most critical oncogenic drivers in LUAD, will be helpful in the development of novel strategies for targeted therapies in lung cancer.

In summary, we have demonstrated that SETD7 inhibits NSCLC progression by downregulating KRAS signaling cascades. SETD7 can methylate KRAS at K182 and K184, further enhancing KRAS ubiquitination and degradation via the recruitment of the E3 ligase, RabGEF1 (**Figure 7**). Our data suggest that the combination of SETD7 and RabGEF1 will be a useful index for prognostic prediction in NSCLC cases.

**Figure 7.**
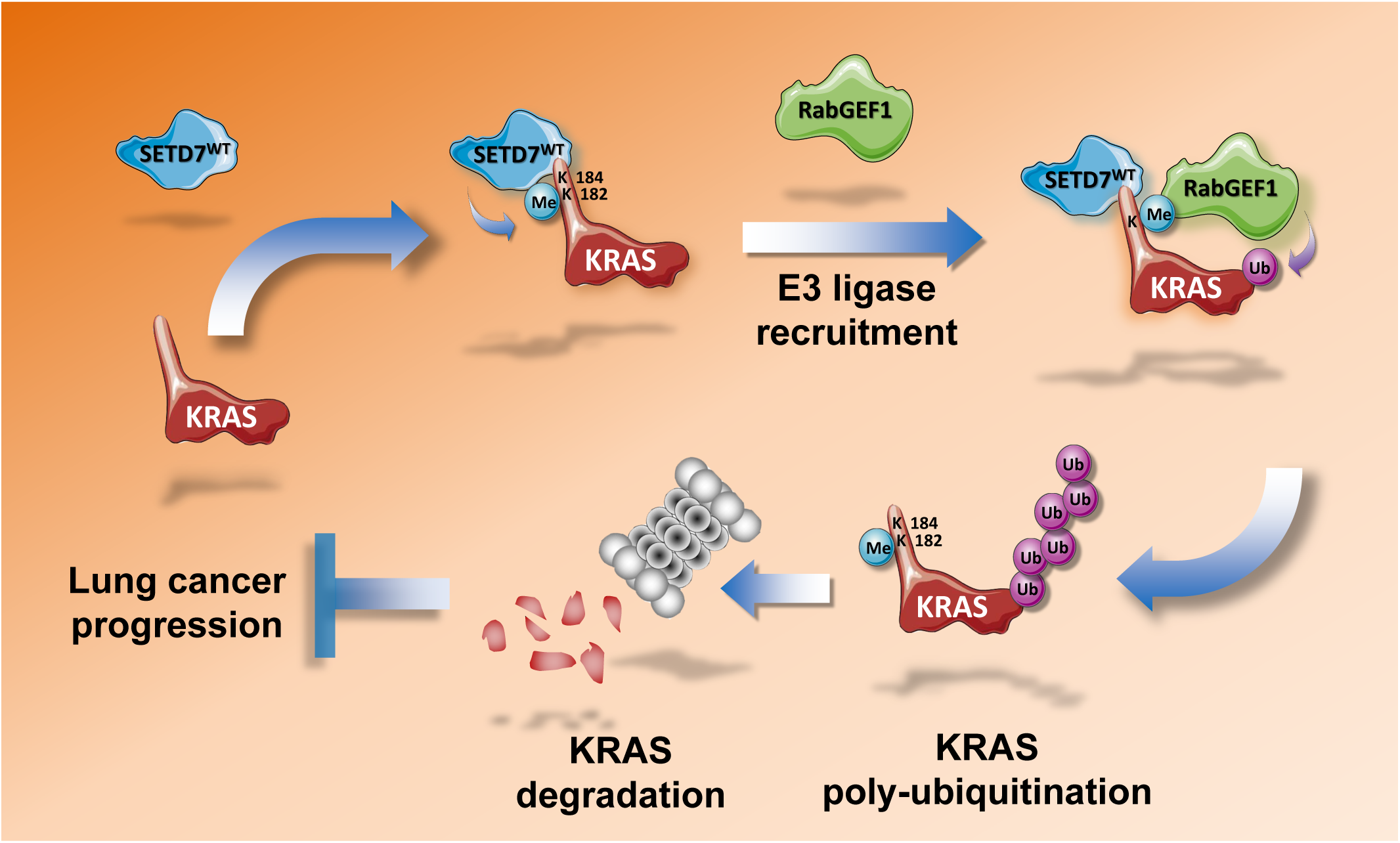
Proposed mechanism of SETD7-mediated KRAS methylation and degradation in lung cancer. SETD7 interacts with KRAS and methylates KRAS at K182 and K184. Then, the E3 ligase, RABGEF1, is recruited in a methylation-dependent manner and promotes polyubiquitination and degradation of KRAS. A reduction in KRAS protein levels suppresses the downstream MEK/ERK and PI3K/AKT cascades and inhibits tumor progression. In lung adenocarcinoma, SETD7 is frequently downregulated, which abolishes its inhibitory effect on KRAS and emancipates the oncogenic role of KRAS.

## Materials and Methods

### Human Lung Cancer Specimen Analysis

Tissue Microarray (TMA) for survival analysis was obtained from 127 patients with lung cancer who underwent potentially curative pulmonary resection at the Department of Thoracic Surgery, the Second Xiangya Hospital of Central South University from 2002-2012. Patients enrolled in this study were based on pathologic diagnosis of lung adenocarcinoma, each sample containing a minimum of 70% tumor cells as determined by study pathologists. Approval was given in advance by our institutional review board, and all patients gave written informed consent. These patients had been subjected to routine staging and definitive surgical resection of the lung and systematic mediastinal lymph node dissection. No patients had been previously treated with chemotherapy and radiotherapy at the time of original operation. Complete clinical record and follow-up data were available for all patients. Overall survival time was calculated from the data of diagnosis to the date of death or the data last known alive.

To examine the SETD7 expression profile, the TMA sections were labeled with an SETD7 antibody (Abcam, 1:200). Immunohistochemical (IHC) staining of TMA sections were scored independently by qualified pathologists blinded to the clinicopathological data, at ×200 magnification light microscopy. Staining intensity was classified as 0 (negative), 1 (weak), 2 (moderate) or 3 (strong). According to these scores, an optimal cut-off point was identified as follows: a staining index score of 0-1 was used to define tumors with low expression and 2-3 indicated high expression of SETD7 protein. Agreement between the evaluators was 95%, and all scoring discrepancies were resolved through discussion between the evaluators. Kaplan-Meier analysis was performed for overall survival.

### Reagents, enzymes, and antibodies

Chemical reagents and reaction buffers were obtained from Sigma-Aldrich (St. Louis, MO, USA). Q5^®^ High-Fidelity DNA Polymerase, DNA ligase and the restriction enzymes were purchased from New England BioLabs (NEB, Ipswich, MA, USA). A list of antibodies used is provided in the Supplementary Materials.

### Construction of plasmids and mutagenesis

Full-length human SETD7 cDNA sequences were amplified from a human mRNA pool generated by RT-PCR using SuperScript II RNase H Reverse Transcriptase (Life Technologies,CA, USA). The SETD7 cDNA was cloned into the *EcoRI/BamHI* site of a pCDH-CMV-MCS-EF1α -Puro vector (System Biosciences, Palo Alto, CA, USA). The sequences were confirmed by Sanger sequencing. Point mutations were performed using Q5 DNA polymerase to generate KRAS K182/184M mutants. The primary DNA template was digested with DpnI for 2 hours at 37°C, then the mutant pCDH-KRAS was transformed and amplified by DH5α competent cells. The primers used for cloning and mutagenesis are provided in the Supplementary Materials.

### Cell culture and stable pools selection

HEK293T cells were obtained from Cell Bank of Chinese Academy of Sciences and cultured in Dulbecco’s modified Eagle’s medium (DMEM; Life Technologies) with 10% fetal bovine serum (FBS; PAN Seratech, Aidenbach, Germany) and 100 U/ml penicillin-streptomycin (Life Technologies). The human non-small-cell lung cancer (NSCLC) cell lines A549, NCI-H358, NCI-H1437, and NCI-H522 were obtained from the American Type Culture Collection (ATCC) and cultured in RPMI1640 (Life Technologies) containing 10% FBS (PAN Seratech) and 100 U/mL penicillin-streptomycin (Life Technologies). HEK293T, A549, NCI-H358, NCI-H1437, NCI-H522, and stable pools were incubated at 37°C with 5% CO_2_. All cell lines were authenticated by short tandem repeat (STR) profiling (Guangdong Huaxi Forensic Physical Evidence Judicial Appraisal Institute, Guangdong, China). The cell lines were passaged fewer than ten times or 6 months after initial revival from frozen stocks; they tested negative for mycoplasma.

For generation of stable pools with SETD7 overexpression or knockdown, NSCLC cell lines were infected by lentivirus expression of SETD7 or SETD7 shRNA and selected by culturing in RPMI1640 complete medium containing 10 μg/mL puromycin (Selleck, TX, USA) for 1 week. SETD7 expression was evaluated by qPCR and western blotting. Functional assays were performed after confirmation of gene expression. The shRNA oligonucleotides used are listed in the Supplementary Materials.

### Real-time PCR

Total RNA was extracted using Trizol reagent (Life Technologies) and cDNA was synthesized using RevertAid First Strand cDNA Synthesis Kit (Thermo Scientific, Waltham, MA, USA) according to the manufacturer’s instructions. All real-time PCR reactions were performed using the CFX Connect Real-Time PCR Detection System (Bio-Rad, Hercules, CA, USA) and the amplifications were carried out using the ChamQ Universal SYBR qPCR Master Mix (Vazyme, Nanjing, China). Each real-time PCR reaction was repeated three times, and target genes were normalized using GAPDH as an internal reference. Primers for real-time PCR were listed in Supplementary Materials.

### Immunoprecipitation and western blot

For immunoprecipitation, HEK293T cells transfected with indicated plasmids were cultured in a 100-mm dish. Cells were collected using IP lysis buffer when they were 80% confluent. Following clarification, the supernatant fractions were used for immunoprecipitation with anti-HA, anti-Flag, or anti-myc antibody and rotated at 5 rpm at 4°C overnight. Next, protein A/G magnetic beads (MedChemExpress, Monmouth Junction, NJ, USA) was added and incubated for 4 h. After washing the magnetic beads several times with binding buffer (25 mM Tris-HCl, pH 7.5,100 mM NaCl, 0.1% NP-40), the immunoprecipitant was finally eluted with sample buffer containing 1% SDS.

Western blotting was performed following immunoprecipitation. The protein concentration was determined using a BCA Protein Assay Kit (Beyotime Biotechnology, Shanghai, China). All protein samples were separated by sodium dodecyl sulfate-polyacrylamide gel electrophoresis (SDS-PAGE), transferred to polyvinylidene fluoride (PVDF) membranes (Merck Millipore, Burlington, MA, USA), hybridized with the corresponding antibodies, and detected using Pierce ECL Western blotting substrate (Thermo Scientific).

### Immunofluorescence

The transfected cells were grown on cover slide and fixed by 50% methanol for 10 min at 4°C. After incubation with anti-SETD7 or anti-KRAS antibodies (1:100) in PBS containing 5% bovine serum albumin over night at 4°C, samples were rinsed with PBS, incubated with Alexa Fluor 488 conjugated anti-mouse or Alexa Fluor 546 conjugated anti-rabbit secondary antibodies (1:300, Life Technologies) in PBS containing 5% bovine serum albumin for 2 hours at room temperature avoiding light. After washing in PBS, samples were mounted by ProLong™ Gold Antifade Mountant with DAPI (Life Technologies) for observation. Images were acquired using a confocal microscopy (LSM880, Carl Zeiss, Jena, Germany).

### Mass spectrometry (MS) analysis of KRAS-interacting proteins

The beads samples obtained from immunoprecipitation experiment were washed three times with pre-cooled PBS buffer to remove the remaining detergent. Then beads samples were incubated in the reaction buffer (1% SDC/100 mM Tris-HCl, pH 8.5/10 mM TCEP/40 mM CAA) at 95 °C for 10 min for protein denaturation, cysteine reduction and alkylation. The eluates were diluted with equal volume of H_2_O and subjected to trypsin digestion overnight by adding 1 μg of trypsin at 37 °C. The peptide was purified using self-made SDB desalting columns. The eluate was vacuum dried and stored at −20 °C for later use.

Mass spectrometry was performed using TripleTOF 5600+ LC-MS/MS system (SCIEX). The peptide sample was inhaled by an autosampler and bound to a C18 capture column (5 μm, 5 × 0.3 mm), followed by elution to the analytical column (300 μm × 150 mm, 3 μm particle size, 120 Å pore size, Eksigent) to carry out the separation. Two mobile phases (mobile phase A: 3% DMSO, 97% H2O, 0.1% formic acid and mobile phase B: 3% DMSO, 97% ACN, 0.1% formic acid) were used to establish the analytical gradient. The flow rate of the liquid phase was set to 5 μL/min. For mass spectrometry IDA mode analysis, each scan cycle contains a full MS scan (m/z range 350-1250, ion accumulation time 250ms), followed by 40 MS/MS scans (m/z range is 100-1500, ion accumulation time 50 ms). The conditions for MS/MS acquisition are set to a parent ion signal greater than 120 cps and a charge number of +2 to +5. The exclusion time for ion repeat acquisition is set to 18 s.

Raw data from TripleTOF 5600 were analyzed with ProteinPilot (V4.5) using the Paragon database search algorithm and the integrated false discovery rate (FDR) analysis function. Spectra files were searched against the UniProt Human reference proteome database using the following parameters: Sample Type, Identification; Cys Alkylation, Iodoacetamide; Digestion, Trypsin; Search Effort was set to Rapid ID. Search results were filtered with unused ≥ 1.3. Decoy hits and contaminant proteins were removed; the remaining identifications were used for further analysis.

### MS analysis of KRAS PTMs

The bands corresponding to the KRAS proteins were cut and subjected to in-gel digestion. Briefly, gels were destained with 50% 50mM NH_4_HCO_3_/50%ACN, were dehydrated with 100%ACN. After reduction and alkylation, proteins were digested with addition of trypsin or chymotrypsin at 37□ overnight. Peptides were dissolved in 0.1%FA and analyzed with LC-MS. An ultra-high pressure nano-flow chromatography system (Elute UHPLC, Buker) was coupled with TimsTOF Pro (Bruker). Liquid chromatography was performed on a reversed-phase column (40cm x 75 um i.d.) at 50 °C packed with Magic C18 AQ 3-μm 200-Å resin with a pulled emitter tip. A solution is 0.1%FA in H2O, and B solution is 0.1%FA in ACN. In 120-min experiments, peptides were separated with a linear gradient from 0%-5% B within 5min, followed by an increase to 30% B within 105min and further to 35% B within 5min, followed by a washing step at 95% B and re-equilibration. The timsTOF pro was operated in PASEF mode. The ddaPASEF files were searched against Swissprot human (downloaded in September, 2018) appendant with common contaminants using Peaks software. The search parameters were set as followed: parent monoisotopic tolerance 15 ppm, product ion tolerance 0.05 Da, modification Carbamidomethyl (C), potential modification protein N-terminal acetylation, methylation on KR, oxidation on M and phosphorylation on STY, and maximum missed cleavage sites 2. The MS/MS spectra of identified modified peptides were validated manually.

### Cell growth and 2-D clonogenic assay

For cell growth assay, cells were seeded in 96-well plates at concentrations of 1-2 × 10^3^ cells per well. Cells were incubated at 37°C in a final volume of 100 µL culture medium per well. Cell viability was evaluated at 0, 24, 48, and 72□hours after cell attachment. Cell Counting Kit-8 (CCK-8, Meliun Biotechnology, China) reagent (10 µL) was added to each well, and the plates were incubated for 2 hours at 37°C. The absorbance was measured at 450 nm.

For 2-D clonogenic assay, cells were then seeded in 6-well plates at a concentration of 500 cells per well. The cells were incubated at 37°C in a final volume of 4 mL culture medium per well. After incubation for 8 to 10 days, the 6-well plates were washed with 1×PBS and the cells were fixed by adding 100% methanol and incubating them at 4°C for 10 minutes. The cells were then stained with 0.5% crystal violet for 10 minutes at room temperature before being washed twice in double-distilled (dd) H_2_O.

### Soft agar assay and 3-D spheroid formation assay

For soft agar assay, cells were seeded in 0.4% low-melting agarose (Sigma-Aldrich) in 6-well plate at a concentration of 5×10^3^ cells per well. The cells were incubated at 37°C for 2-3 weeks, then stained with 0.005% crystal violet for 1 hour at room temperature before being destained in ddH_2_O.

In spheroid formation assay, cells were seeded in 96-well microplate (U-bottom) with cell-repellent surface (Greiner Bio-One GmbH, Frickenhausen, Germany) at a concentration of 2×10^3^ cells per well. Cells were incubated at 37°C in a final volume of 200 µL culture medium per well for 2 weeks. The volume of spheroids were monitored by inverted microscopy and calculated by use of the modified ellipsoid formula 1/2(length × width^2^).

### Computer simulation of the molecular structure

For the computer simulation, SETD7 and KRAS crystal structures were obtained from the Protein Data Bank (PDB)(Berman et al., 2000). The G117-K366 sequence of SETD7 (PDB ID: 1O9S)(Xiao et al., 2003) and the M1-C185 sequence (PDB ID: 6CCX)(Fang et al., 2018) of KRAS, both determined by X-ray diffraction, were selected as the initial docking structures. The HADDOCK server (version 2.4, http://milou.science.uu.nl/services/HADDOCK2.2/), set to default parameter values, was used to perform simulations of KRAS docking with the SET7 binding pocket. The most common structures from clusters 1, 2, 3, and 5 were fed into the dynamic molecular simulation. In this study, all simulations were performed using the AMBER package. The FF14SB force field was applied for the protein preparation simulation. The complex was immersed in TIP3PBOX. Counter ions were added to neutralize the system. First, the entire system was minimized until convergence occurred. After minimization, the system, with restraint on all heavy atoms, was gradually heated up to 300 K, in NVT ensemble. The force constant of restraint was set to 100 kcal/(mol·rad2). Then, the system was relaxed in NPT ensemble in four stages, where the force constant was set to 50, 20, 10, and 5 kcal/(mol·rad2), respectively. The entire simulation lasted for 100 ns, with a snapshot taken every ps. Cluster analysis was performed on the last 20-ns trajectory. The linkage method was selected and the cutoff was set to 4.0 Å.

### Xenografts in nude mice

All animal research procedures were performed according to the detailed rules of the Animal Care and Use Ethics Committee of Shenzhen University Health Science Center, and all animals were treated in strict accordance with protocols approved by the Institutional Animal Use Committee of the Health Science Center, Shenzhen University. Male *nu*/*nu* nude mice, 4 to 6 weeks old, were subcutaneously injected in their back with 5×10^6^ stable pool cells containing A549-pCDH-puro, A549-pCDH-SETD7, H1437-scramble control, and H1437-pLKO.1-shSETD7#1 or H1437-pLKO.1-shSETD7#2. Tumor volume was monitored once per week. Tumor volume was calculated according to the formula: volume = length × width^2^ / 2. After 4 to 7 weeks, mice were euthanized, and their tumor xenografts were harvested for weighing and following IHC staining for SETD7, KRAS, and Ki-67.

### Statistical analyses

All *in vitro* experiments were performed at least in triplicate. The results of each experiment are presented as the mean ± standard deviation. Data analyses were performed using Microsoft Excel 2010 Professional Plus (Version 14.0.7237.5000) and Prism 6 (Version 6.01). Two-tailed, unpaired Student’s t-tests were used to compare differences between two groups with similar variances. For all tests, a p-value <0.05 was considered to indicate a statistically significant difference.

## Acknowledgments

This study was supported by grants from the National Natural Science Foundation of China (81974366, 81773019), the Natural Science Foundation of Guangdong Province (2019A1515010210, 2020A1515011408), Guangdong Provincial Science and Technology Program (2019B030301009), the Shenzhen Municipal Government of China (JCYJ20180507182427559, GJHZ20180418190559891), and Shenzhen Key Medical Discipline Construction Fund (SZXK060). Plasmid pCMV-myc-ubiquitin was a gift from Prof. Xingzhi Xu; plasmid pvxl-flag-RabGEF1 was a gift from Dr. Xifeng and plasmid pCDH-puro-KRAS^G12D^ was a gift from Dr. Hongbin Ji. The authors would like to thank Dr. Xi Chen (SpecAlly Life Technology Co., Ltd, Wuhan, China) for mass spectrometry data analysis, Chengli Weng for animal experimental support, the Instrumental Analysis Center of Shenzhen University (Lihu Campus) for the technical support, and Dr. Jessica Tamanini of ETediting for editing the manuscript prior to submission.

## Author contributions

T.X. and C.C. conceived and designed the experiments. C.C. carried out most of the experiments and analyzed the data. M.Z. helped with the construction of plasmids and lentivirus packaging. T.X. helped with the animal studies. S.F. and H.Z. prepared tissue microarray and performed IHC staining and scoring of SEDT7. C.Z. performed the LC-MS/MS and proteomic analysis. J.Z. performed the computer simulation of SETD7 and KRAS crystal structures. S.L. and W.G. contributed to the planning and discussion of the project. D.Z. supervised the entire project. T.X. and C.C. wrote the manuscript, designed the layout of figures and tables with contributions from all of the authors.

## Supplementary Figure Legends

**Figure S1.**
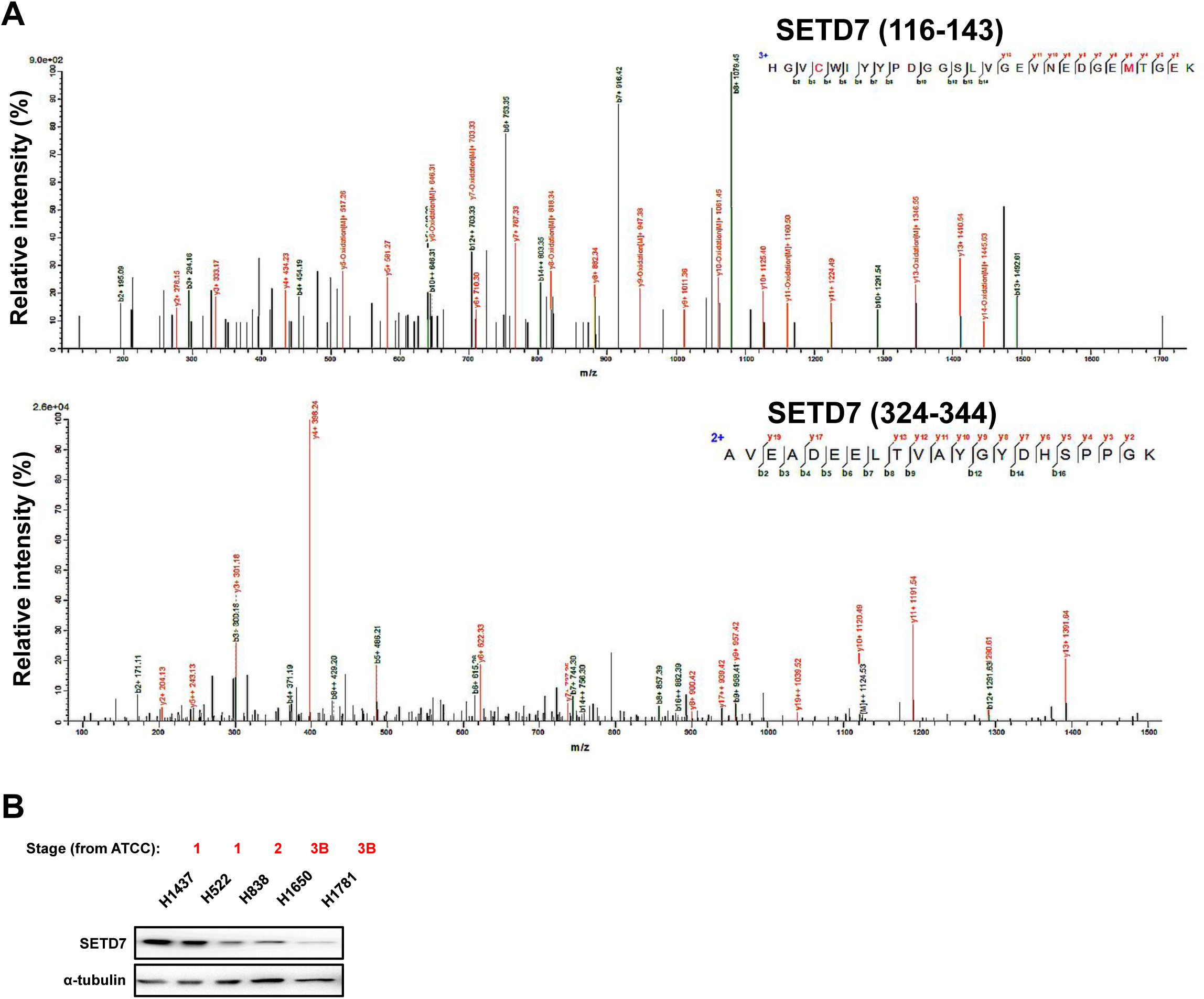
Correlation between SETD7 expression and tumor progression in non-small-cell lung cancer (NSCLC) cell lines and clinical specimens. **(A)** Two representative MS2 spectra of SETD7 peptides (116-143 and 324-344) from 3×HA-KRAS group. **(B)** Western blotting (WB) analysis of SETD7 expression in NSCLC cell lines from the American Type Culture Collection (ATCC). Staging information is shown above each cell line in red font.

**Figure S2.**
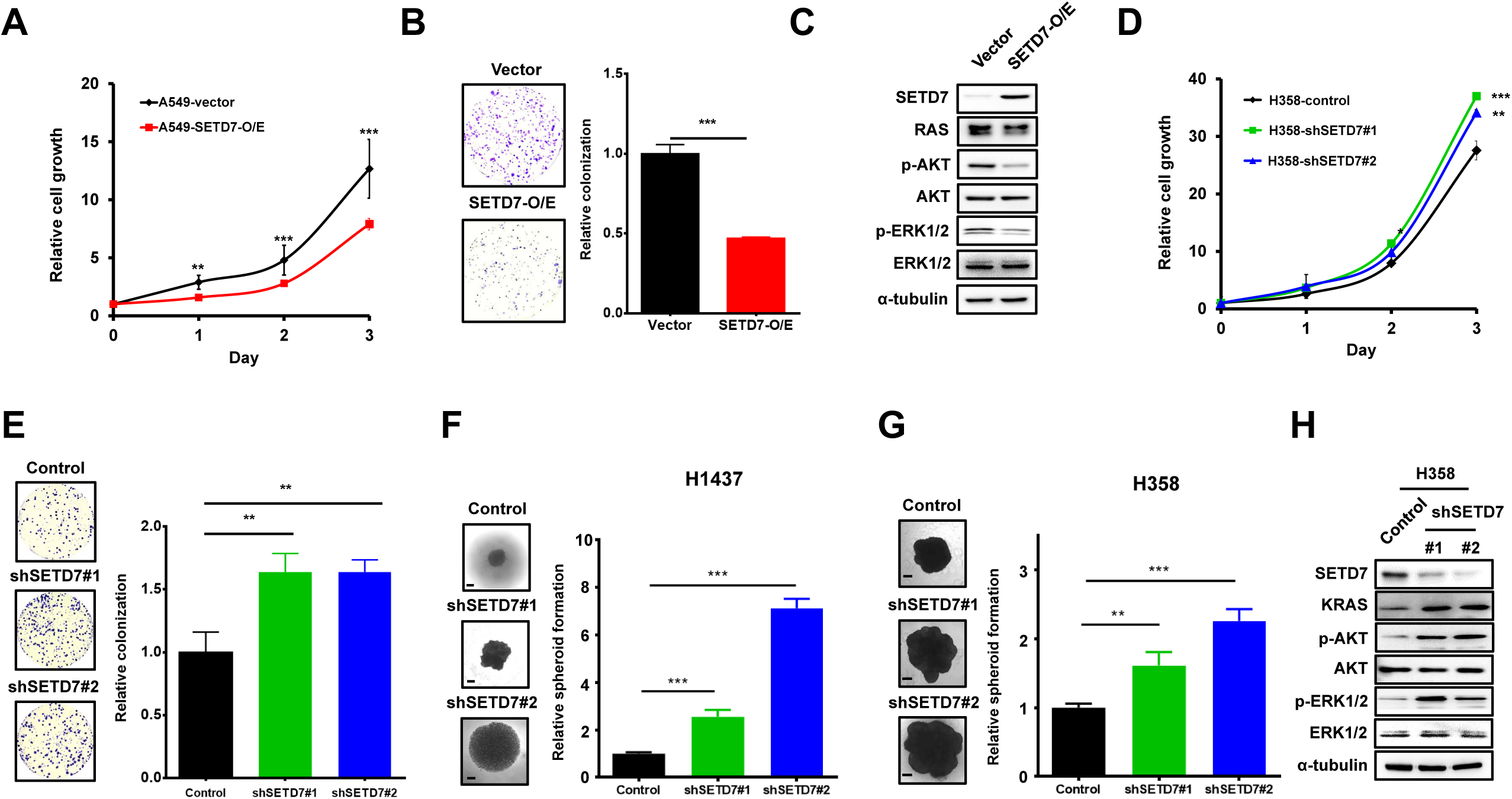
The tumor-suppressive role of SETD7 in NSCLC cell lines. **(A-B)** Cell proliferation (A) and two-dimensional (2-D) colony formation (B) of A549-vector and A549-SETD7-OE stable cells was measured separately. **(C)** WB analysis of SETD7 and RAS-related signaling pathways in A549 stable cells. **(D-E)** Cell proliferation (D), 2-D colony formation (E) of H358-control and two SETD7-KD stable cells were measured separately. **(F-G)** 3-D spheroid growth of H1437 (F) and H358 (G) controls and two SETD7-KD stable cells was measured separately. **(H)** WB analysis of SETD7 and RAS-related signaling pathways in H358 stable cells. \**P*<0.05; \*\**P*<0.01; \*\*\**P*<0.001.

**Figure S3.**
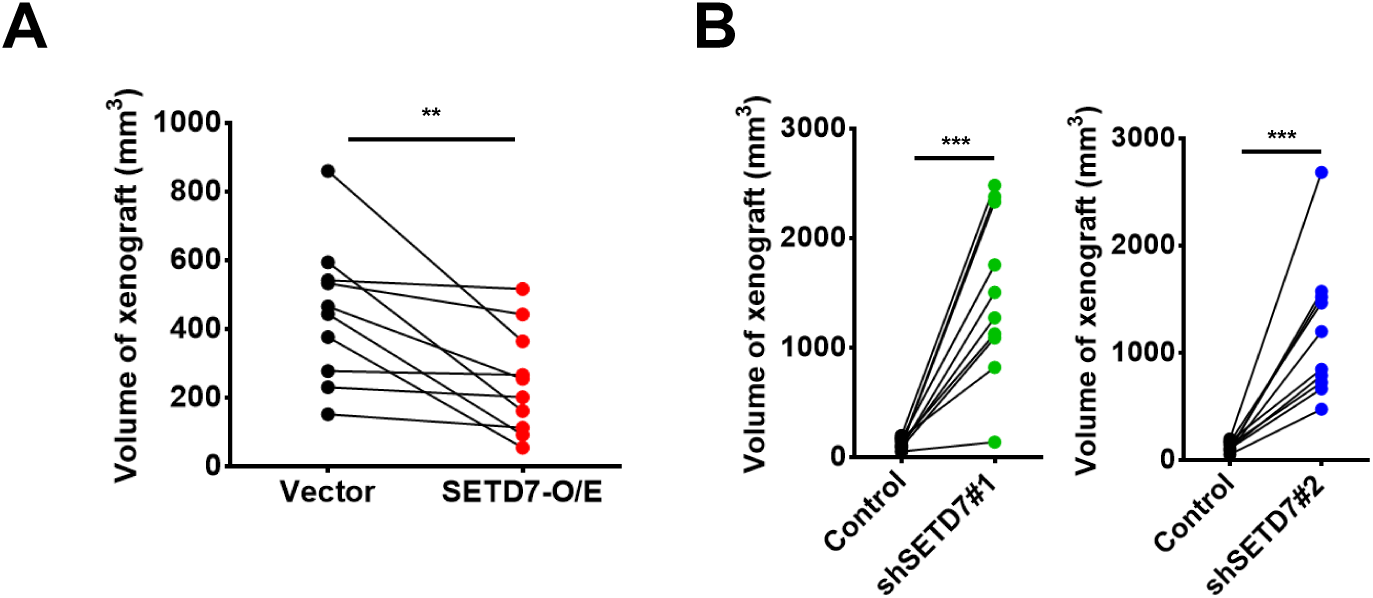
The anti-cancer effect of SETD7 in a xenograft assay. **(A)** Tumor volumes of xenografts derived from A549 vector and SETD7-OE stable pools were measured 7 weeks after tumor cell injection. \*\**P*<0.01. **(B)** Tumor volumes of xenografts derived from H1437 vector and two SETD7-KD stable pools were measured 4 weeks after tumor cell injection. \*\*\**P*<0.001.

**Figure S4.**
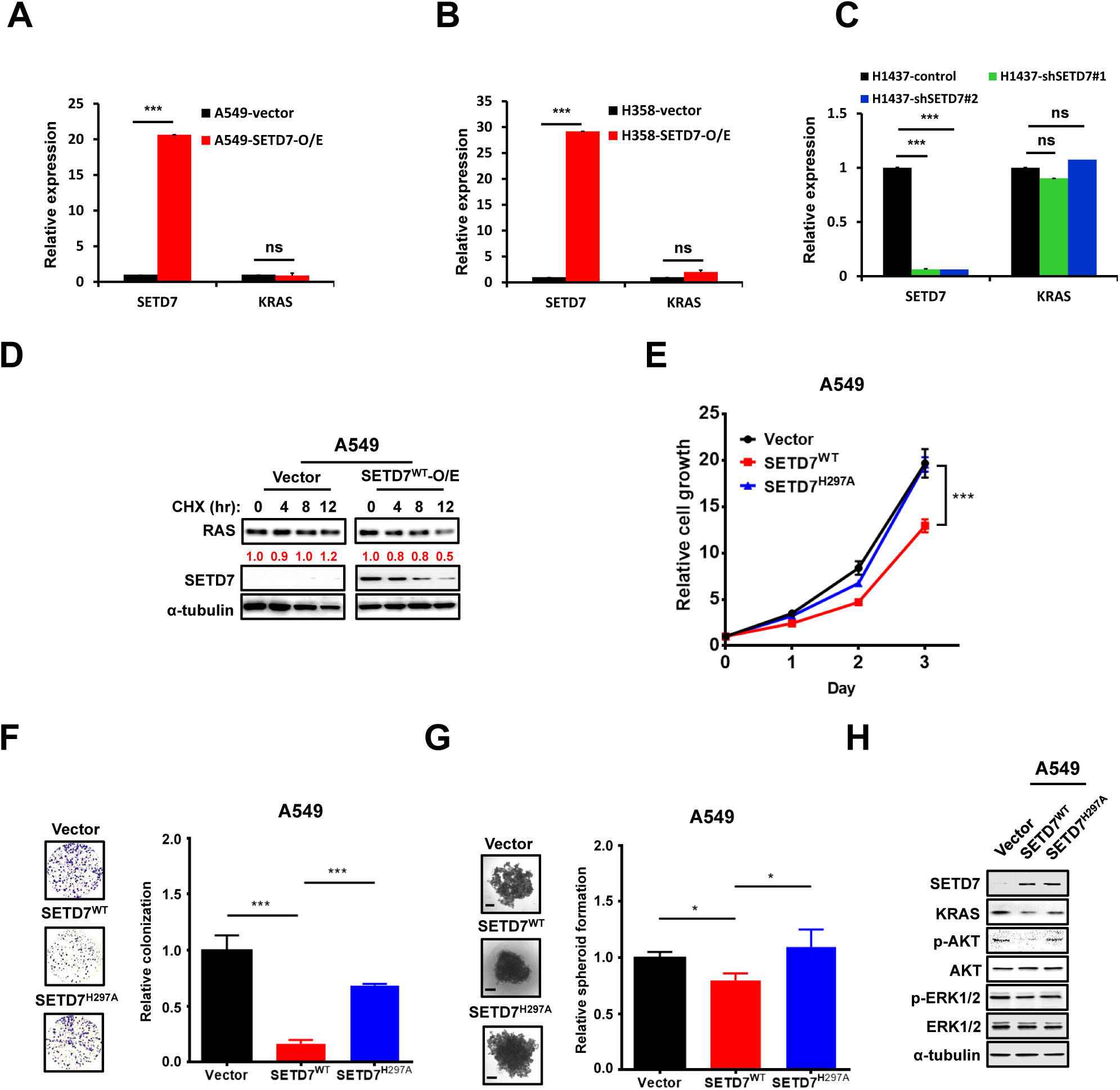
The anti-cancer effect of SETD7 is dependent on its catalytic activity. **(A-C)** KRAS mRNA expression was detected by qPCR in A549 (A) and H358 (B) stable cells with SETD7 overexpression, or H1437 (C) stable cells with SETD7 knockdown. **(D)** WB analysis of A549-vector and A549-SETD7-O/E stable cells with CHX (100 μg/mL) treatment, as indicated. The relative RAS protein level is labeled in red. **(E)** Cell proliferation of A549 vector, SETD7^WT^, and SETD7^H297A^ stable cells was measured using a CCK-8 assay. **(F-G)** Two-dimensional (2-D) colony formation (F) and 3-D spheroid growth (G) of A549 vector, SETD7^WT^, and SETD7 ^H297A^ stable cells were measured separately. The bar graphs represent the quantification of numbers of colonies on plates or spheroids in dishes with a cell-repellent surface, from three independent experiments. **(H)** WB analysis of SETD7 and RAS-related signaling pathways in A549 stable cells. \**P*<0.05, \*\*\**P*<0.001. ns, not significant.

**Figure S5.**
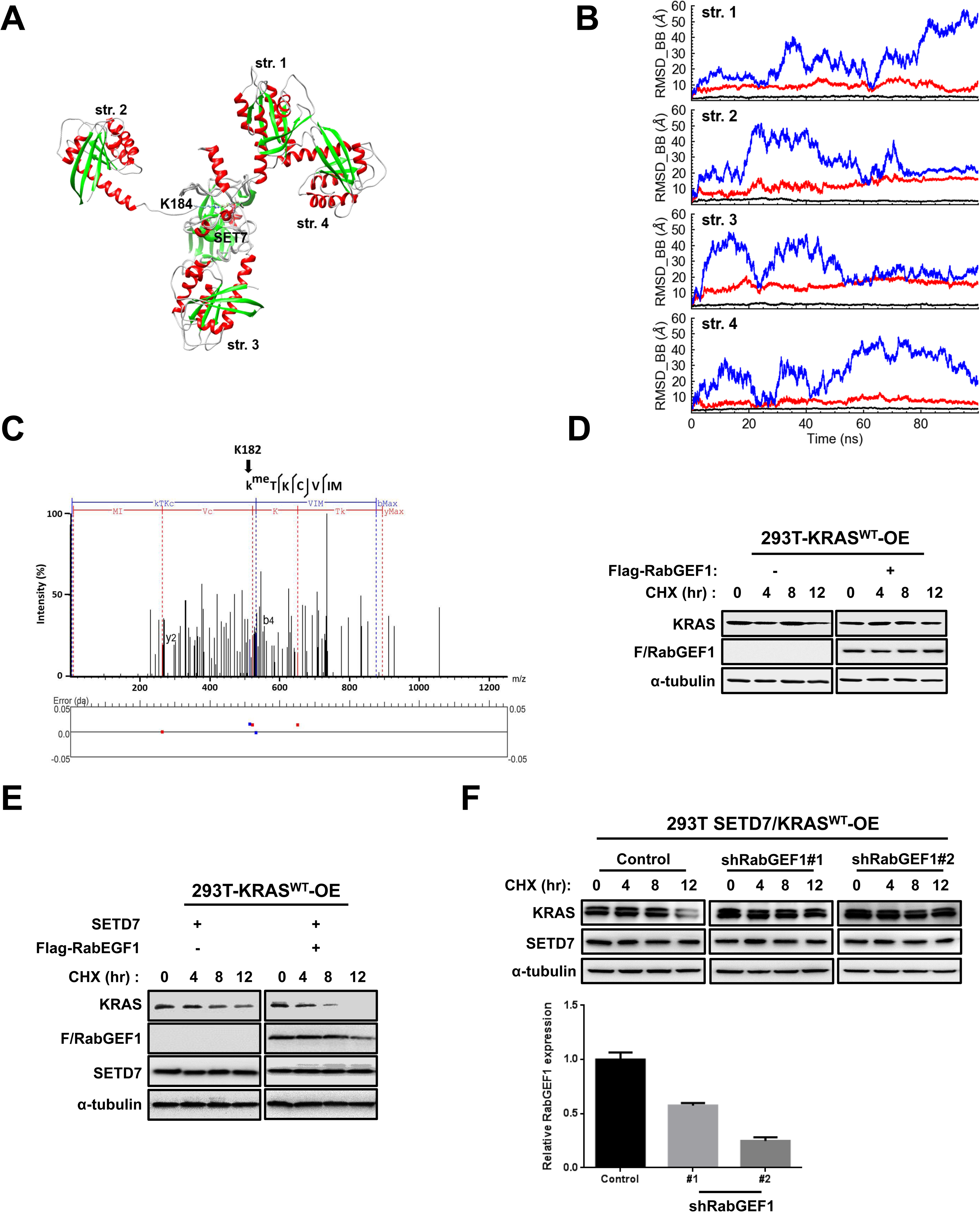
The interaction between SETD7 and KRAS. **(A)** A computer simulation of the interaction between SETD7 and KRAS. The SETD7 sequence G117-K366 was applied to perform the molecular dynamics (MD) simulation. Binding sites of the SETD7 SET domain (PDB ID: 1O9S): TYR245, GLY264, ASN265, THR266, LEU267, SER268, HIS293, ALA295, TYR305, TYR335, GLY336, TYR337 (245, 264, 265, 266, 267, 268, 293, 295, 305, 335, 336, 337); Prediction of the methylation site of KRAS (PDB ID: 5OCG): K182 or K184 (181, 182, 183, 184, 185). The listed structures from HADDOCK were selected as the initial structures for MD simulations. **(B)** The root-mean-square deviation (RMSD) calculated for the backbone atoms evolved with the simulation time. **(C)** KRAS peptide modified with methylation on lysine 182 residue was identified by LC-MS/MS. **(D)** WB analysis of 293T cells transfected with KRAS and RABGEF1 or KRAS alone with CHX (100 μg/mL) treatment, as indicated. **(E)** WB analysis of 293T cells co-transfected with KRAS and SETD7 with or without RabGEF1, following indicated CHX (100 μg/mL) treatment. **(F)** WB analysis of 293T cells co-transfected with KRAS and SETD7, and control or RabGEF1 shRNAs, following indicated CHX (100 μg/mL) treatment (upper panel). Knockdown efficiency of RabGEF1 was monitored by qPCR (lower panel).

**Figure S6.**
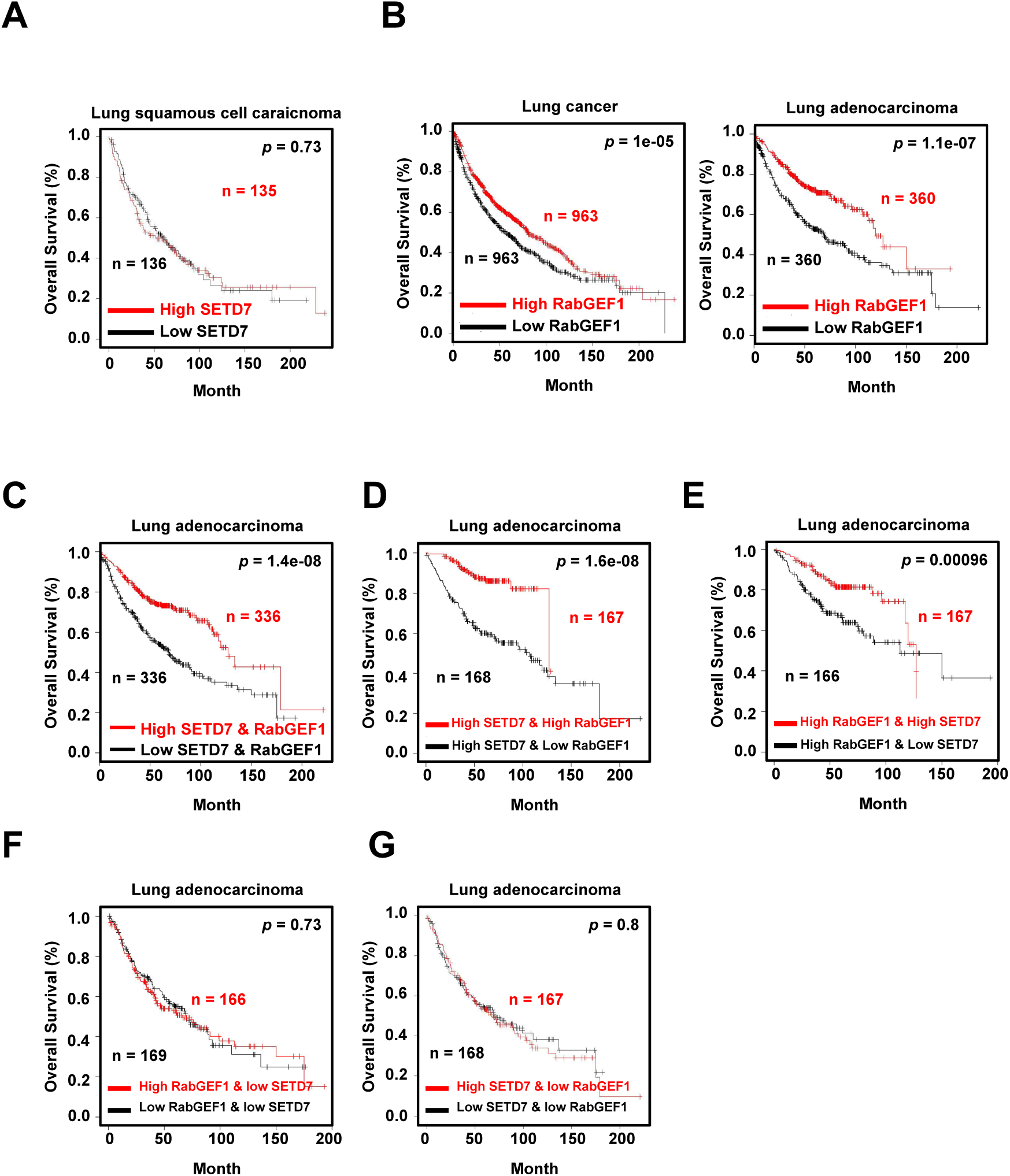
Kaplan-Meier plotter survival analysis for SETD7 and RabGEF1 mRNA levels in lung cancer and lung adenocarcinoma (LUAD) **(A)** KM plotter survival analysis for SETD7 mRNA in patients with lung squamous cell carcinoma (n = 271). **(B)** KM plotter survival analysis for RABGEF1 mRNA levels (probe set, 218310_at) in lung cancer (n = 1925) and LUAD (n = 720) patient samples. **(C-G)** KM plotter survival analysis for the combination of SETD7 and RABGEF1 mRNA levels in patients with LUAD. The comparisons were performed between patients with both high SETD7 and RabGEF1 expression (n = 336) and patients with both low SETD7 and RabGEF1 expression (n = 336) (C); Patients with high SETD7 expression were divided into two groups according to whether their RABGEF1 expression was high (n = 167) or low (n = 168) (D); Patients with high RABGEF1 expression were divided into two groups according to SETD7 expression, high (n = 167) or low (n = 166) (E); Patients with low SETD7 expression were divided into two groups according to RABGEF1 expression, high (n = 166) or low (n = 169) (F); Patients with low RABGEF1 expression were divided into two groups according to SETD7 expression, high (n = 167) or low (n = 168) (G). In the above KM plotter survival analyses, medium was used as a cutoff to divide the samples into low and high groups.

