## Supplementary Material and Methods for "Methylation of KRAS at lysine 182 and 184 by SETD7 promotes KRAS degradation"

**Supplementary Materials and Methods**

**Antibodies list**

| Antibody | Cat. no. | Supplier | Dilution factor |
| --- | --- | --- | --- |
| HA-tag | sc-7392 | Santa Cruz Biotechnology( Dallas, TX, USA) | 1:2000 |
| Flag-tag | F1804 | Sigma-Aldrich (St. Louis, MO, USA) | 1:2000 |
| β-actin | CW0096A | CoWin BioSciences (Cambridge, MA, USA) | 1:2000 |
| α-tubulin | 11224-1-AP | ProteinTech Group, Inc. (Rosemont, IL, USA) | 1:2000 |
| Myc-tag | 2276 | Cell Signaling Techonlogy (Danvers, MA, USA) | 1:1000 |
| phospho-Akt | 4060 | Cell Signaling Techonlogy (Danvers, MA, USA) | 1:1000 |
| Akt | 9272 | Cell Signaling Techonlogy (Danvers, MA, USA) | 1:1000 |
| phospho-Erk1/2 | 4070 | Cell Signaling Techonlogy (Danvers, MA, USA) | 1:1000 |
| Erk1/2 | 4695 | Cell Signaling Techonlogy (Danvers, MA, USA) | 1:1000 |
| SETD7 | ab124708 | Abcam (Cambridge, England) | 1:2000 |
| RAS | ab52939 | Abcam (Cambridge, England) | 1:2000 |
| KRAS | WH0003845M1 | Sigma-Aldrich (St. Louis, MO, USA) | 1:2000 |
| Ki67 | GB13030-2 | Servicebio (Wuhan, China) | 1:200 |
| Histone 3 | 17168-1-AP | ProteinTech (St. Rosemont, IL, USA) | 1:1000 |
| mono-methyl-lysine | ab23366 | Abcam (Cambridge, England) | 1:2000 |

**Oligo-nucleutide list**

| **Cloning primers** | | | |
| --- | --- | --- | --- |
| SETD7- Forward | 5’-ATTTGAATTCATGGATAGCGACGAC-3’ | | |
| SETD7- Reverse | 5’-TCCTCTAGATTATCAGGCGTAGTCG-3’ | | |
| KRAS  K182/184M-Forward | 5’-AAAAGAAGAAAAAGAAGTCAATGACAATGTGTGTAATTATGTAA-3’ | | |
| KRAS  K182/184M-Reverse | 5’-TTACATAATTACACACATTGTCATTGACTTCTTTTTCTTCTTTT-3’ | | |
| SETD7-H297A Forward | 5’-TGCCTCCTTGGGACACAAGGCAAATGCCTCCTTCACTCCAAACTGCATCTACG-3’ | | |
| SETD7-H297A-Reverse | 5’-CGTAGATGCAGTTTGGAGTGAAGGAGGCATTTGCCTTGTGTCCCAAGGAGGCA-3’ | | |
| **qPCR primers** | | | |
| SETD7-Forward | | 5’-TATGTCCACTGAAGAAGG-3’ | |
| SETD7-Reverse | | 5’-AAGAAGAGCATTGGTAGA-3’ | |
| KRAS-Forward | | 5’-AAGTAGTAATTGATGGAGAA-3’ | |
| KRAS-Rverse | | 5’-CTCTATAATGGTGAATATCTTC-3’ | |
| GAPDH-Forward | | 5’-AGGTGAAGGTCGGAGTCAAC-3’ | |
| GAPDH-Reverse | | 5’-AGTTGAGGTCAATGAAGGGG-3’ | |
| **Oligo-nucleutide for knockdown** | | | |
| shSETD7-#1-Top | | | 5’-CCGGGGGAGTTTACACTTACGAAGACTCGAGTCTTCGTAAGTGTAAACTCCCTTTTTG-3’ |
| shSETD7-#1-Bottom | | | 5’-AAAAGGGAGTTTACACTTACGAAGACTCGAGTCTTCGTAAGTGTAAACTCCC-3’ |
| shSETD7-#2-Top | | | 5’-CCGGGGACCGCACTTTATGGGAAATCTCGAGATTTCCCATAAAGTGCGGTCCTTTTTG-3’ |
| shSETD7-#2-Bottom | | | 5’-AATTCAAAAAGGACCGCACTTTATGGGAAATCTCGAGATTTCCCATAAAGTGCGGTCC-3’ |
| shRABGEF1-#1-Top | | | 5’-CCGGGGATGCAAACTCGTGGGAAAGCTCGAGCTTTCCCACGAGTTTGCATCCTTTTTG-3’ |
| shRABGEF1-#1-Bottom | | | 5’-AATTCAAAAAGGATGCAAACTCGTGGGAAAGCTCGAGCTTTCCCACGAGTTTGCATCC-3’ |
| shRABGEF1-#2-Top | | | 5’-CCGGGGCGATCACAGATATCATTGACTCGAGTCAATGATATCTGTGATCGCCTTTTTG-3’ |
| shRABGEF1-#2-Bottom | | | 5’-AATTCAAAAAGGCGATCACAGATATCATTGACTCGAGTCAATGATATCTGTGATCGCC-3’ |
